# Automated Collective Variable Discovery for MFSD2A transporter from molecular dynamics simulations

**DOI:** 10.1101/2024.04.19.590308

**Authors:** Myongin Oh, Margarida Rosa, Hengyi Xie, George Khelashvili

## Abstract

Biomolecules often exhibit complex free energy landscapes in which long-lived metastable states are separated by large energy barriers. Overcoming these barriers to robustly sample transitions between the metastable states with classical molecular dynamics (MD) simulations presents a challenge. To circumvent this issue, collective variable (CV)-based enhanced sampling MD approaches are often employed. Traditional CV selection relies on intuition and prior knowledge of the system. This approach introduces bias, which can lead to incomplete mechanistic insights. Thus, automated CV detection is desired to gain a deeper understanding of the system/process. Analysis of MD data with various machine learning algorithms, such as Principal Component Analysis (PCA), Support Vector Machine (SVM), and Linear Discriminant Analysis (LDA)-based approaches have been implemented for automated CV detection. However, their performance has not been systematically evaluated on structurally and mechanistically complex biological systems. Here, we applied these methods to MD simulations of the MFSD2A (Major Facilitator Superfamily Domain 2A) lysolipid transporter in multiple functionally relevant metastable states with the goal of identifying optimal CVs that would structurally discriminate these states. Specific emphasis was on the automated detection and interpretive power of LDA-based CVs. We found that LDA methods, which included a novel gradient descent-based multiclass harmonic variant, termed GDHLDA, we developed here, outperform PCA in class separation, exhibiting remarkable consistency in extracting CVs critical for distinguishing metastable states. Furthermore, the identified CVs included features previously associated with conformational transitions in MFSD2A. Specifically, conformational shifts in transmembrane helix 7 and in residue Y294 on this helix emerged as critical features discriminating the metastable states in MFSD2A. This highlights the effectiveness of LDA-based approaches in automatically extracting from MD trajectories CVs of functional relevance that can be used to drive biased MD simulations to efficiently sample conformational transitions in the molecular system.

**STATEMENT OF SIGNIFICANCE:** To elucidate the biological mechanisms of pertinent biomolecules, it is crucial to understand their complex free energy landscapes. Such landscapes are often constructed from molecular dynamics (MD) simulations using collective variable (CV)-guided enhanced sampling methods. Identifying proper CVs for this task is critical but can be challenging with traditional intuition-based approaches. Here we propose an automated protocol for CV discovery which is based on linear discriminant analysis (LDA) for dimensionality reduction of MD data. By applying the protocol to MD simulations of the MFSD2A lysolipid transporter, a structurally and mechanistically complex biological system, we show that LDA-based methods efficiently detect system-specific CVs that accurately classify different metastable states of MFSD2A and are highly interpretable in a detailed structural context.

## INTRODUCTION

Biomolecules are highly dynamic and flexible. They manifest high-dimensional complex free energy landscapes characterized by a hierarchy of energy barriers which involve biomolecular motions that take place on multiple timescales^1-4^. Biologically important functions are often associated with conformational transitions between metastable states. To unravel the structure-dynamics-function relationship of biomolecules, it is thus imperative to investigate the diffusive behavior of the system on the free energy hypersurface at an atomistic level. Computational molecular dynamics (MD) simulations are a very powerful and valuable tool for this purpose as it can provide microscopic view of the biologically relevant processes. However, the free energy surfaces governing biomolecular dynamics often contain multiple metastable states separated by large energy barriers, creating kinetic bottlenecks^5^. This limitation restricts the scope and timescale that can be explored with MD simulations, as the presence of the barriers introduces significant statistical errors in measurement of structural, thermodynamic, and kinetic properties of biologically relevant transitions when sampled during MD simulations in an unbiased manner^6^.

To address the sampling issue, enhanced sampling techniques based on collective variables (CVs) have become standard practice^6-16^. CVs are any differentiable functions of atomic coordinates that provide a low-dimensional projection of the conformational space of a biomolecular system without missing important information about the system. After projection, they must retain the ability to distinguish important metastable states and transition states, and all slow motions must be captured using only a minimal number of CVs along which to drive slow transitions when biased^6-7,10^. Good CVs provide lower variance estimates of properties of interest, enable the definition of effective bias potentials for enhanced sampling, and aid in visualizing the free energy landscape^6^.

For structurally and mechanistically complex biomolecular systems, it is highly challenging to discover good CVs based purely on physical intuition. However, the CV detection process has been revolutionized and automated with the surge in data availability and recent advancements in the development of algorithms that integrate various machine learning (ML) techniques into MD data analysis^6-7^. Among these, linear dimensionality reduction (DR) algorithms stand out owing to their ease of implementation, direct interpretability, and reduced computational demands. Examples of such approaches include principal component analysis (PCA), used to identify a subspace that most preserves the configurational variance within a molecular simulation trajectory,^6,17^ and time-lagged independent component analysis (tICA), used to identify slow CVs that span a subspace that preserves the maximum kinetic content by maximizing the autocorrelation function^6,18^. While both PCA and tICA are unsupervised linear transformation techniques (i.e., they do not require data labeling), they suffer from different types of limitations: in PCA it is assumed that the large amplitude motions the algorithm detects are related to slow motions involved in transitions between biologically relevant metastable states^19-21^; a significant drawback of using tICA for CV design is its requirement for either unbiased or biased sampling of slow transitions through MD simulations^15^.

Linear discriminant analysis (LDA) is another class of linear DR algorithms commonly used for classification tasks. In contrast to PCA and tICA, LDA operates on labeled data which is projected onto a subspace where the separation of different classes is maximized^6, 22-23^. This maximization is achieved by optimizing the ratio of a between-class variance to a within-class variance. Importantly, all the basis vectors of the linear transformation of the data in LDA (i.e., CVs) can be identified directly from unbiased sampling of local fluctuations within each metastable state *without the need to sample transitions between these states*. Here, we apply LDA to MD simulations of MFSD2A (Major Facilitator Superfamily Domain 2A), a Na^+^-dependent omega-3 fatty acid membrane transporter which is enriched at the blood-brain barrier where it plays a central role in a number of neurological processes^24-28^. We choose MFSD2A as our case study due to its mechanistic complexity, and because it is a structurally and functionally well-characterized system^29-32^. Indeed, our recent studies^29-30^ combining MD simulations with cryo-electron microscopy (cryo-EM) and transport assay experiments have revealed multiple functionally relevant metastable states in MFSD2A: outward-facing (OFS) and inward-facing (IFS) states determined from the cryo-EM as well as an intermediate occluded state (OcS), predicted from our large-scale MD simulations (**Figure 1**). Our objective here was to apply the LDA methodology to unbiased atomistic MD simulations of MFSD2A in these three structurally distinct metastable states in order to uncover CVs that would allow for structural classification of OFS, OcS, and IFS. Such CVs can be useful for future design of enhanced sampling MD simulations to elucidate detailed molecular mechanisms of state-to-state transitions in MFSD2A.

**Figure 1.**
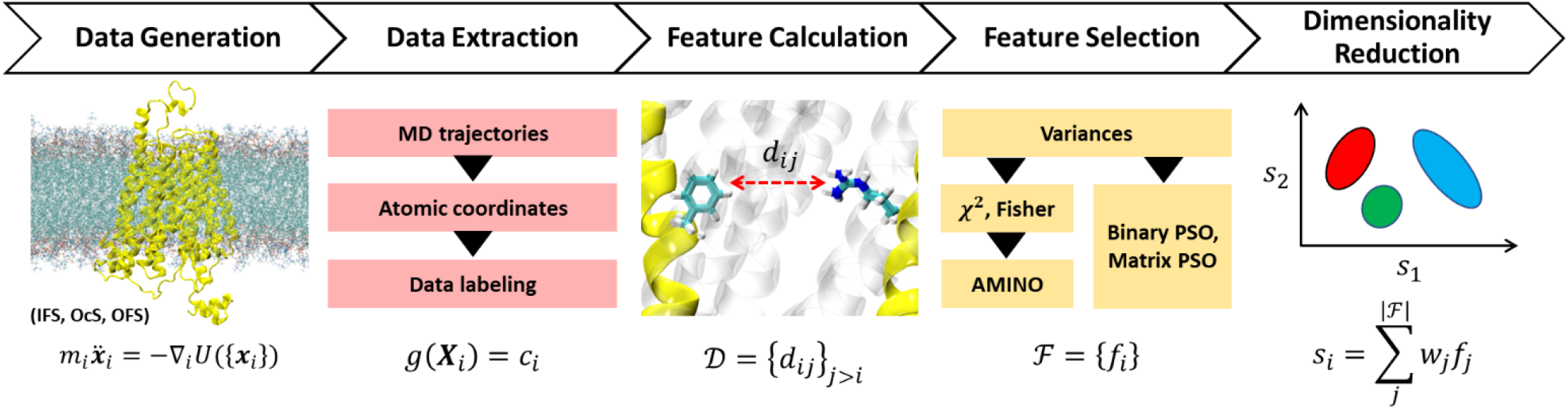
Schematic workflow of our protocol for designing CVs for MFSD2A conformational transitions. The process begins with unbiased MD simulations for each metastable state or class (IFS, OcS, and OFS), followed by extraction of atomic coordinates **x**_*i*_ and label assignment *g* for each conformation **X**_i_. Here, the label c_i_ is one of the three classes. Pairwise distances d_ij_ among residues (using either center-of-mass distances or the C_α_-C_α_ distances) are calculated and compiled into a distance set 𝒟. From this, the most informative features f_j_ are selected to form the final feature set *ℱ*. CVs (s_i_) are then generated using linear DR algorithms as a linear combination of the features f_j_ with w_j_ as the corresponding weights.

To systematically evaluate the performance of LDA for automated CV detection, we considered various formulations of the approach (see Methods), such as Fisher LDA (FLDA)^33^, and harmonic variants of LDA (HLDA) proposed by Zheng et al (ZHLDA)^34^ and Mendel et al (MHLDA)^23^. In addition, we present here and test a new LDA variant, the gradient descent-based multiclass HLDA (GDHLDA) method. GDHLDA leverages the strengths of ZHLDA and MHLDA by integrating the harmonic mean approaches for both within-class and between-class variances and relies on gradient descent on the Stiefel manifold to identify the orthonormal basis vectors for the linear transformation (see Methods). We extend our analysis to a comparative evaluation of CVs derived from other linear and nonlinear machine-learning approaches, including PCA^17^, support vector machines (SVM),^35^ and deep convolutional neural networks (DCNNs)^36^. We show that the LDA-based DR methods enable the generation of system-specific CVs that clearly discriminate the metastable states of MFSD2A and therefore can be instrumental for the future design of enhanced sampling MD simulations to elucidate the detailed molecular mechanisms underlying state-to-state transitions of MFSD2A. Specifically, our analysis identified conformational shifts in the transmembrane helix 7 (TM7) of MFSD2A and in residue Y294 on the intracellular side of this helix as one of the major determinants in the structural classification of the OFS, OcS, and IFS. Overall, we found that LDA-based methods, including GDHLDA, outperformed PCA in class separation, exhibiting remarkable consistency in extracting features critical for distinguishing the metastable states of MFSD2A. Furthermore, the features extracted by LDA methods were consistent with those obtained from our previously developed non-linear DCNN technique for pattern recognition from MD data. With that, this work places an emphasis on the automated detection and interpretive power of LDA-based CVs.

## METHODS

### CV Design Protocol

Our protocol for designing CVs for MFSD2A conformational transitions comprises five main steps (see **Figure 1** for the complete workflow of the protocol and refer to the subsequent sections below for a detailed explanation of each step in the workflow). First, we generate data directly from unbiased MD simulations for each metastable state. Second, we extract atomic coordinates from these simulations, assigning each conformation to a class based on the set of criteria established in our previous work^29-30^. Third, we compute pairwise distances among residues (using either distances between their center-of-masses or *C*_*α*_-*C*_*α*_ distances), forming a comprehensive distance set. The fourth step focuses on feature reduction and consists of multiple stages: we initially remove constant (within fluctuations) and quasi-constant features based on their variances. To further select the most informative features, we implement filter and wrapper methods. Our filter approach encompasses the χ^2^ test^37^ and Fisher scores^38-39^, which are then followed by the Automatic Mutual Information Noise Omission (AMINO)^40^ process, specifically designed to remove noisy and redundant features. For the wrapper methods, we utilize binary particle swarm optimization (BPSO)^34^ and matrix PSO (MPSO). Once the feature selection is complete, we apply different linear DR algorithms, including PCA, FLDA, ZHLDA, MHLDA, and GDHLDA, to construct CVs as a linear combination of the selected features.

### Molecular Constructs for atomistic MD simulations

Atomistic MD simulations were carried out for MFSD2A in three distinct conformational states: OFS, OcS, and IFS. For the OFS system, we used the previously established^30^ homology model for full-length *Gallus gallus* (ggMFSD2A) based on the structural template derived from the cryo-EM structure (PDB: 7N98)^31^ of *Mus musculus* (mmMFSD2A). Similarly, for the IFS, the structural template was obtained from the cryo-EM structure (PDB: 7MJS)^29^ of ggMFSD2A, as used in the earlier computational studies^29^. In both OFS and the IFS models, the transporter is in *apo* (ligand-free) state.

The OcS model was acquired from our previous simulation work^30^. Specifically, from the 1.6 μs-long MD trajectory that exhibited an OFS-to-OcS transition during the first 600 ns, we analyzed the last 4000 frames, corresponding to 640 ns, as reported in^30^. During this trajectory, the LPC-18:1 (lysophosphatidylcholine 18:1) substrate from the extracellular leaflet spontaneously permeated the central region of MFSD2A, where it engaged in Na^+^-bridged interactions with E312^30^. This interaction facilitated the protein’s transition into the OcS conformation. In order to select a representative frame of the OcS conformation as a starting structure for the new set of simulations reported in the current work, we used multiple criteria established previously^30^: we considered cut-off distances from the LPC-18:1 substrate headgroup phosphate atoms (< 4.8Å) and Na^+^ ions (< 2.6Å) to E312’s carboxyl carbon, combined with the analysis of the root-mean-square deviation (RMSD) measure calculated for the residues comprising the 12 TMs of MFSD2A (residue IDs 39 to 68, 75 to 101, 110 to 129, 137 to 167, 171 to 202, 233 to 263, 292 to 318, 328 to 353, 358 to 374, 382 to 416, 424 to 450, and 465 to 491). The RMSD variance of these 4000 frames was below 1 Å, indicating their high similarity. Among these, we selected a structure, using GROMACS gmx cluster tool^41^, that exhibited the RMSD value closest to the centroid to serve as our representative conformation. The final choice of the OcS structure met all three criteria: LPC and Na^+^ coordination with residue E312, and closest RMSD to the centroid of structures.

Frames for both the IFS and OFS constructs were selected from the previously reported simulations^29-30^ as the final snapshots from one of the independent replicates (as described^30^, the OFS model was simulated in 48 replicates with cumulative sampling time of 74 μs ^30^, and the IFS model was run in 12 replicates with cumulative sampling time of 11.4 μs^29^. This selection ensured that the membrane-embedded and solvated protein systems (as described in ^29,30^) were well-equilibrated prior to using them as starting structures for the new simulations reported here. Notably, the selected IFS frame exhibited a 1.065 Å RMSD compared to IFS cryoEM (PDB: 7MJS), while the selected OFS frame displayed a 1.201 Å RMSD compared to OFS cryoEM (PDB: 7N98), indicating minimal changes in overall protein conformation during the previous MD. For more detailed specifications of the MFSD2A system in the three states, please see our previous publications^29,30^. Briefly, the ggMFSD2A OFS and IFS models were embedded in a homogeneous membrane with 483 and 484 POPC lipids, respectively, and the OcS was inserted into a heterogeneous membrane consisting of 484 POPC and 15 LPC-18:1 lipids. Each protein-membrane complex was surrounded by a solution box containing TIP3P water molecule models and Na^+^ and Cl^−^ ions to maintain a 0.15 M ionic concentration. The simulated systems each contained ∼230,000 atoms, including explicit hydrogens^29,30^.

### MD Simulations

Since the above membrane-embedded and solvated/ionized systems were previously well-equilibrated^29,30^, here, to initialize new simulations from these systems, velocities for all atoms were randomly regenerated after which the constructs underwent extensive multi-replicate MD simulations. In total, 36 independent 500 ns-long replicates (12 per each conformational state), were conducted, resulting in cumulative sampling of ∼18 *μs*. On these timescales, fluctuations of the protein around the three states (OFS, OcS, and IFS) are expected to be sufficiently sampled. The MD simulations were performed with OpenMM 7.4^42^. Particle Mesh Ewald (PME)^43-44^ was implemented for electrostatic interactions. The runs were performed at 310 K and under 1 bar isothermal-isobaric (NPT) ensemble conditions using semi-isotropic pressure coupling, and with 4 fs integration time-step (with hydrogen mass repartitioning)^45^. MonteCarloMembraneBarostat and Langevin thermostats were used to maintain constant pressure and temperature, respectively. Additional parameters for these runs included: “friction” set to 1.0/ps, “EwaldErrorTolerance” 0.0005, “rigidwater” True, and “ConstraintTolerance” 0.000001. The van der Waals interactions were calculated applying a cut-off distance of 12 Å and switching the potential from 10 Å. For all simulations we used the latest CHARMM36m force-field for proteins and lipids,^46^ as well as recently revised CHARMM36 force-field for ions which includes non-bonded fix (NBFIX) parameters.^47^ For all the standard MD data analyses and visualization VMD software version 1.9.4 was used.^48^

### Feature Calculation and Selection

In our approach, we opt for pairwise distances as molecular features to define CVs. We focus on two distinct sets of pairwise distances: 𝒟_RE_, involving distances between the geometric centers of the residues, and 𝒟_CA_, involving distances between the *C*_*α*_ atoms. We considered only the residue pairs in the TM helices 1-12 (see above). For feature selection, we identified constant (within fluctuations) and quasi-constant features by calculating the variance of each feature and discarding those from 𝒟 whose variance fell below a predetermined threshold. The thresholds were determined by identifying the elbow point, where the variance plateau begins, using the Kneedle algorithm^49^ on the variance plot (**Figure S1a,b**). Consequently, the thresholds were set at 3.19 Å^2^ for 𝒟_RE_ and 1.75 Å^2^ for 𝒟_CA_.

To trim the feature set further towards identification of the most informative features, we tested five distinct criteria: (1) the χ^2^ test statistic, (2) the Fisher score, (3) BPSO, (4) MPSO with loose regularization (MSPO-L) and (5) MPSO with strong regularization (MPSO-S). The first two criteria were implemented using Scikit-learn^50^, a well-established Python module for machine learning, while BPSO and MPSO were executed with NiaPy^51^, a versatile Python microframework designed for simple implementation of nature-inspired optimization algorithms.

(1) The χ^2^ test statistic between each feature and the target was calculated as:

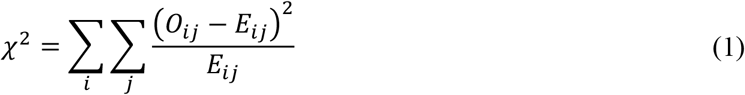

where *O*_*ij*_ and *E*_*ij*_ are the observed and expected frequencies, respectively, summed across the categories^36^. Formally, the χ^2^ test is strictly applicable to the analysis of categorical variables. As our dataset comprised continuous variables, a transformation to the categorical variables was needed first. To this end, we employed the quantile-based discretization function from the widely used pandas library (qcut function)^52^ to categorize our continuous variables into discrete bins. Specifically, the range of each continuous variable was segmented into five equal-sized quantiles, and for each trajectory frame the continuous variable was assigned one of the five discrete categories (0, 1, 2, 3, 4) based on the value of the variable in this frame. Our empirical analysis indicated that a five-bin model was optimal, revealing a more extensive set of features compared to models with fewer categories, while still capturing similar key features. Subsequently, we applied the χ^2^ test to the discretized dataset. A high χ^2^ value indicates that the hypothesis of independence is incorrect and thus there is a strong relationship between the feature and the target^37^. We established the threshold of 1460 features for 𝒟_RE_ and 1594 features for 𝒟_CA_, using the Kneedle algorithm to detect the elbow point the χ^2^ test plot (**Figure S1c,d**), similarly to the variance identification method.
(2) The Fisher score^38-39^ *F* evaluates the discrimination power of features by comparing the variance between classes to the variance within classes. This is mathematically represented as

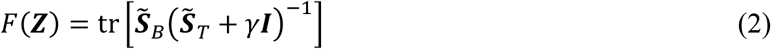

where ***Z*** is the reduced data matrix containing a subset of features, and ***I*** is the identity matrix. Here, 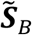 and 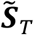 are the between-class and total scatter matrices, respectively, in the reduced data space, defined as

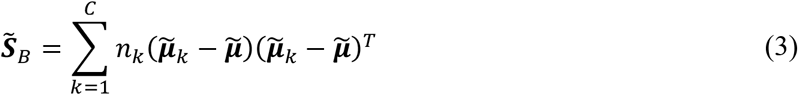

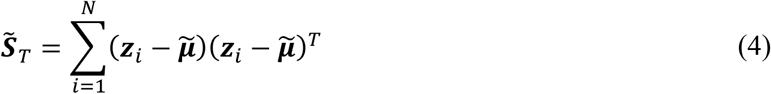

where *C* and *N* denote the total numbers of classes and data points, respectively, *n*_k_ is the number of data points in class *k*, and 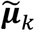 and 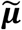 are the mean vectors of class *k* and the overall data, respectively, and *γ* in Eq. (2) is a positive regularization parameter introduced to prevent the singularity of 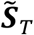. A higher Fisher score suggests that the feature is more effective at distinguishing between classes^38-39^. We set the threshold at 4014 features for 𝒟_RE_ and 3093 features for 𝒟_CA_, determined by identifying the elbow point on the Fisher score plot (**Figure S1e, f**). The χ^2^ test and the Fisher score analyses were followed by AMINO on the remaining feature set to eliminate noisy and redundant features. AMINO utilizes a mutual information-based distance metric to extract a minimally redundant set of features from a larger set, and then uses clustering, combined with rate-distortion theory, to identify representative features from each cluster that maximize information about the system^40^. Specifically, the AMINO algorithm consists of the following key steps: (1) The input features are clustered using a mutual information-based distance metric, which assesses the similarity between every pair of features. This step identifies groups of features that carry similar information. (2) From each cluster, the most representative feature is selected using *k*-medoids clustering, which employs actual data points as centroids, in conjunction with rate-distortion theory. (3) The optimal number of clusters needed to represent the entire set of features that AMINO started with is determined The jump method^53^ is used for estimating the true number of clusters in this step. The resulting features *f*_*i*_ constitute the final feature set ℱ. For this study, we set a maximum of 10 clusters and utilized 30 bins for the analysis.
(3) We used BPSO, a heuristic optimization approach, as another feature selection method^35^. This technique draws inspiration from the social behaviors of biological swarms to efficiently identify optimal feature combinations for classification tasks^35, 54-56^. In BPSO, a swarm of particles represents potential solutions within a multi-dimensional feature space. Each particle’s movement is guided by two key factors: its own best previous experience (cognitive part reflecting personal experience) and the best experience of all the other members (social part reflecting collective experience). Each particle represents a candidate solution, and its status is characterized by its ‘position’ and ‘velocity’. A particle’s position ***X*** in this context represents a specific combination of features, while its velocity ***V*** dictates how it explores the feature space. The optimization process begins by initializing a population of particles with random positions and velocities. Given a total of *N* features, each particle’s position is represented as an *N*-dimensional vector with elements ranging between 0 and 1. These elements are then compared with a threshold of 0.5, resulting in a binary solution vector ***x***, representing a subset of selected features. To evaluate the classification performance of each particle, a bagging classifier comprising an ensemble of 10 base estimators with linear SVMs is trained and tested using the features selected by the particle. The fitness of each particle, determined by the classification accuracy and the size of a feature subset, is then evaluated. In this study, the following fitness function^55^ is used:

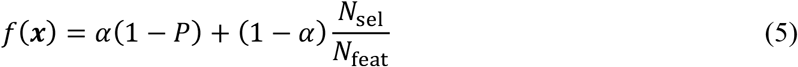

where *P* is the classification accuracy, *N*_sel_ is the number of selected features, *N*_feat_ is the total number of features available, *α* ∈ [0,1] is a weighting factor that balances classifier performance and subset size (*α* was set to 0.99 based on the previous work)^57-59^. After evaluating the fitness of the entire swarm, each particle’s velocity is updated according to the following equation:

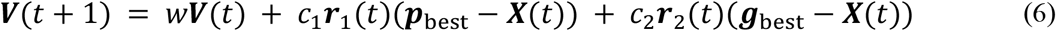

where *t* is the iteration index, ***X***(*t*) is the particle position, ***p***_best_ is the particle’s historical best position, ***g***_best_ is the historical best position of the entire swarm, ***r***_1_(*t*) and ***r***_2_(*t*) are two random vectors in [0, 1] range that share the same dimensionality with the feature space. In Eq. (6), the multiplication between vectors is element-wise (Hadamard product), and *w, c*_1_, *c*_2_ are inertia, cognitive and social acceleration constants, respectively. In this work, we used the default values^60^ for the parameters: *w* = 0.7, *c*_1_ = *c*_2_ = 2.0. The particles’ positions are then updated as ***X***(*t*+ 1) = ***X***(*t*) + ***V***(*t* + 1). This iterative process— comprising feature selection and evaluation using a predefined classification algorithm—continues until a termination criterion, such as a maximum number of iterations or a convergence threshold, is met. In this work, we established the maximum number of iterations for each PSO run to range between 100 and 1000. To avoid computational complexity, we adopt a hierarchical approach; specifically, we conduct multiple runs of BPSO, each time using the reduced feature set obtained from the preceding run as the input. This iterative process is structured to repeat eight to nine times, while maintaining an exceptionally high classification accuracy (> 99%), ultimately aiming to narrow down the feature set to only five selected features. The PSO algorithm is thus terminated once this criterion of feature selection is achieved.
(4-5) To enhance computational efficiency and robustness of the BPSO-SVM classifier, we introduce a MPSO-SVM classifier, which leverages the benefits of signal amplification and noise reduction by averaging concurrent motions. This variant of BPSO employs a ‘position matrix’ ***X*** with size (*k, N*_feat_), where *k* is a user-defined parameter that represents the final dimensionality of the reduced conformational space (i.e., the final number of CVs) (with *k* ≪ *N*_feat_). Each row of the binary solution matrix ***X*** encapsulates a specific combination of selected features, i.e., one CV is created through the average of the *N*_k,sel_ selected features on each row. The linear SVM classifier is then trained and tested in the *k*-dimensional space, using these CVs as the input. The benefit of MPSO-SVM classifier compared to the conventional BPSO-SVM classifier is that it avoids the high-dimensional space, potentially comprising up to *N*_feat_ features, encountered in the BPSO-SVM classifier, making MPSO-SVM particularly beneficial for high-dimension datasets. The fitness function of the MPSO-SVM is defined following the same principle as for the BPSO-SVM:

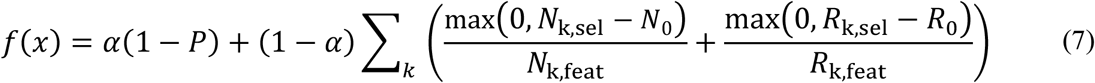

In the above, the first term accounts for the performance, and the second term performs regularizations according to the complexity of the selected features. *N*_k,sel_ and *N*_k,feat_ are the numbers of selected and available features, respectively, in the *k*-th dimension, and *R*_k,sel_ and *R*_k,feat_ are the numbers of related residues to the feature sets. *N*_0_ and *R*_0_ represent the maximum numbers of features and residues allowed in each dimension before any penalty is applied to the regularization term. In this work, we employed either a strong regularization with *N*_0_ = 1 and *R*_0_ = 3, or a loose regularization with *N*_0_ = 5 and *R*_0_ = 6.

### LDA approach for dimensionality reduction

LDA is a well-established DR technique specifically well-suited for pattern recognition and classification problems^61-63^. In practice, LDA is particularly advantageous when dealing with datasets where the number of dimensions is greater than the number of samples, a common scenario in many high-dimensional biological and physical datasets^63^. LDA aims to project high-dimensional data onto a lower-dimensional subspace while maximizing class separability. This is achieved through a linear transformation that maximizes the ratio of between-class variance to the within-class variance in the projected data.

### FLDA formalism

Mathematically, the goal of LDA is to find a linear combination of features that separates classes as much as possible, and in FLDA^33^, this is done by maximizing the following criterion

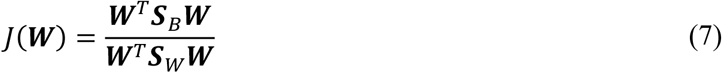

where ***W*** represents the matrix of transformation vectors (projection matrix), ***S***_*B*_ is the between-class scatter matrix, and ***S***_*W*_ is the within-class scatter matrix. These scatter matrices are defined as:

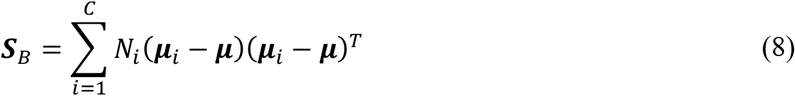

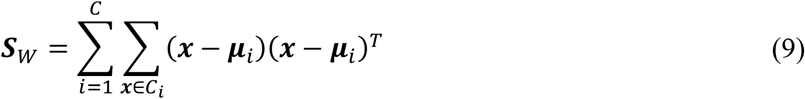

Here, *N*_*i*_ is the number of samples in class *i, μ*_*i*_ is the mean vector of class *i, μ* is the global mean vector of the data, and *C*_*i*_ represents the set of samples belonging to class *i*. The optimal ***W*** is found as the eigenvectors corresponding to the largest eigenvalues 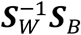 namely,

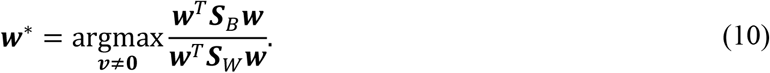

These eigenvectors define the directions (or decision boundaries) in the new subspace that maximizes the class separation. Several mathematical aspects should be emphasized in FLDA. First, ***S***_***W***_ and ***S***_***B***_ are always positive semi-definite, meaning that they symmetric real matrices whose eigenvalues are non-negative. Second, ***S***_***W***_ must have a full rank (*i*.*e*., invertible). Third, ***W*** is a *p* × *k* matrix where *p* = |*ℱ*| is the cardinality of the final feature set and *k* is the reduced dimension subject to *k* ≤min(*C* − 1, *p*). Fourth, the orthogonality of ***W*** is not always guaranteed (*i*.*e*., ***W***^*T*^***W*** ≠ *I*). This is because, despite the symmetry of the scatter matrices, the product 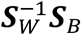 is not necessarily symmetric. Lastly, it is assumed that the data in each class follow a Gaussian distribution with identical covariance matrices, which might not always hold true in real-world data. Nevertheless, many studies attest to its robustness and versatility as a DR tool^64-66^.

### HLDA approaches

The use of arithmetic means in evaluating both ***S***_*W*_ and ***S***_***B***_ matrices are limitations of FLDA as (1) the approach fails to acknowledge the significance of metastable states characterized by smaller fluctuations which are typically more difficult for the system to transition into and out of, and therefore should be weighted more heavily^23^, and (2) the approach treats all the distances equally when the between-class scatter matrix is calculated, overlooking the importance of small between-class distances that, if prioritized, could significantly enhance class separation^34^. To overcome the inherent limitations of FLDA, alternative formulations for CV design that leverage different weighting schemes have been proposed. Zheng et al.^34^ proposed a ZHLDA variant in which the reciprocal of the weighted harmonic mean of pairwise between-class distances is minimized, whereas Mendels et al.^23^ introduced another formulation (MHLDA) which employs the harmonic average of within-class variances. CVs obtained from ZHLDA and MHLDA have been shown to surpass FLDA CVs in separability and interpretability when applied to a range of processes, from the liquid-superionic phase transition of silver iodide^23^ to the folding of chignolin^66^ and the crystallization of Na and Al^67^, as well as in various chemical reactions^22-23,68^.

### GDHLDA formulation

Here, we developed a variant of HLDA, termed GDHLDA (gradient descent-based HLDA) which takes advantage of the strengths of ZHLDA and MHLDA approaches by integrating the harmonic mean approaches for both within-class and between-class variances and, similar to ZHLDA, uses the Stiefel gradient descent algorithm for optimization. Specifically, in GDHLDA a constrained trace ratio optimization problem on the Stiefel manifold is considered (see Algorithm 1), with the objective of finding an orthonormal basis set (or projection matrix) ***W*** that minimizes the objective function *J*, given by:

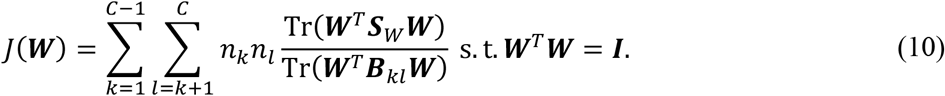

Here, *C* is the number of classes, *n*_*i*_ is the count of data points in class *i*, ***S***_*W*_ is the within-class scatter matrix, and ***B***_*kl*_ is the pairwise between-class scatter matrix, defined as follows:

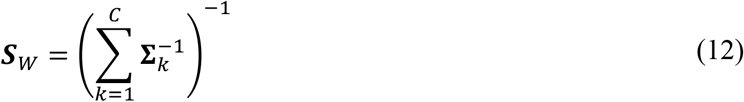

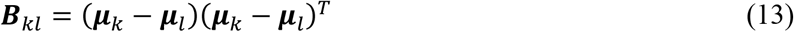

where ∑_*k*_ is the covariance matrix which is defined as

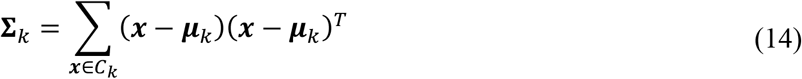

where *C*_*k*_ is class *k* and *μ*_*k*_ is the mean vector of class *k*. As in MHLDA, we utilize the harmonic average for ***S***_*W*_. The constraint ***W***^*T*^***W*** = ***I*** ensures that ***W*** remains on the Stiefel manifold and thus the linear independence of the eigenvectors^34, 69-70^. To preserve this manifold structure, ***W*** is updated as:

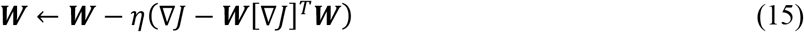

where *η* is the step size^34^. Given that the manifold-preserving gradient enforces the condition ***W***^*T*^***W*** = ***I*** only to the first order, we periodically re-project ***W*** onto the manifold using the singular value decomposition (SVD) every 20 iterations^34^. Stated differently, if SVD(***W***) = ***U***∑***V***^*T*^, then ***W*** is set to ***UV***^*T*^. Furthermore, we incorporate a strict stopping criterion: if ***W*** converges and shows no change over 500 iterations, the algorithm terminates. Notably, GDHLDA circumvents the need for eigendecomposition or matrix inversion and captures all possible orthonormal basis sets effectively (see section “Geometrical Analysis on the Solution Spaces Given by the Linear DR Algorithms” and **Figure S2** in Supplemental Materials). The components of each column vector of ***W*** represent the weights *w*_*i*_ of the features *f*_*i*_ in each CV *s*_*i*_; namely, *s*_*i*_ is expressed as a linear combination of *f*_*i*_ as described by the equation:

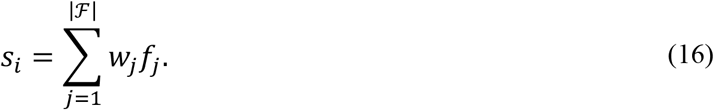

To conduct a comprehensive comparative analysis, we built CVs using a suite of the following linear algorithms: PCA, FLDA, MHLDA, ZHLDA, and GDHLDA. Principal Components (PCs) were calculated with the Scikit-learn^50^ Python module. The LDA-based CVs – FLDA, MHLDA, and ZHLDA – were generated using our Python scripts written in-house (https://github.com/KhelashviliLab/Automated-CV-Design). Specifically, the MHLDA CVs were constructed using the protocol detailed in the work of Mendels et al.^23^, while the ZHLDA CVs were prepared following the methodology outlined by Zheng et al.^34^.

#### Algorithm 1.

Stiefel Gradient Descent Algorithm for HLDA

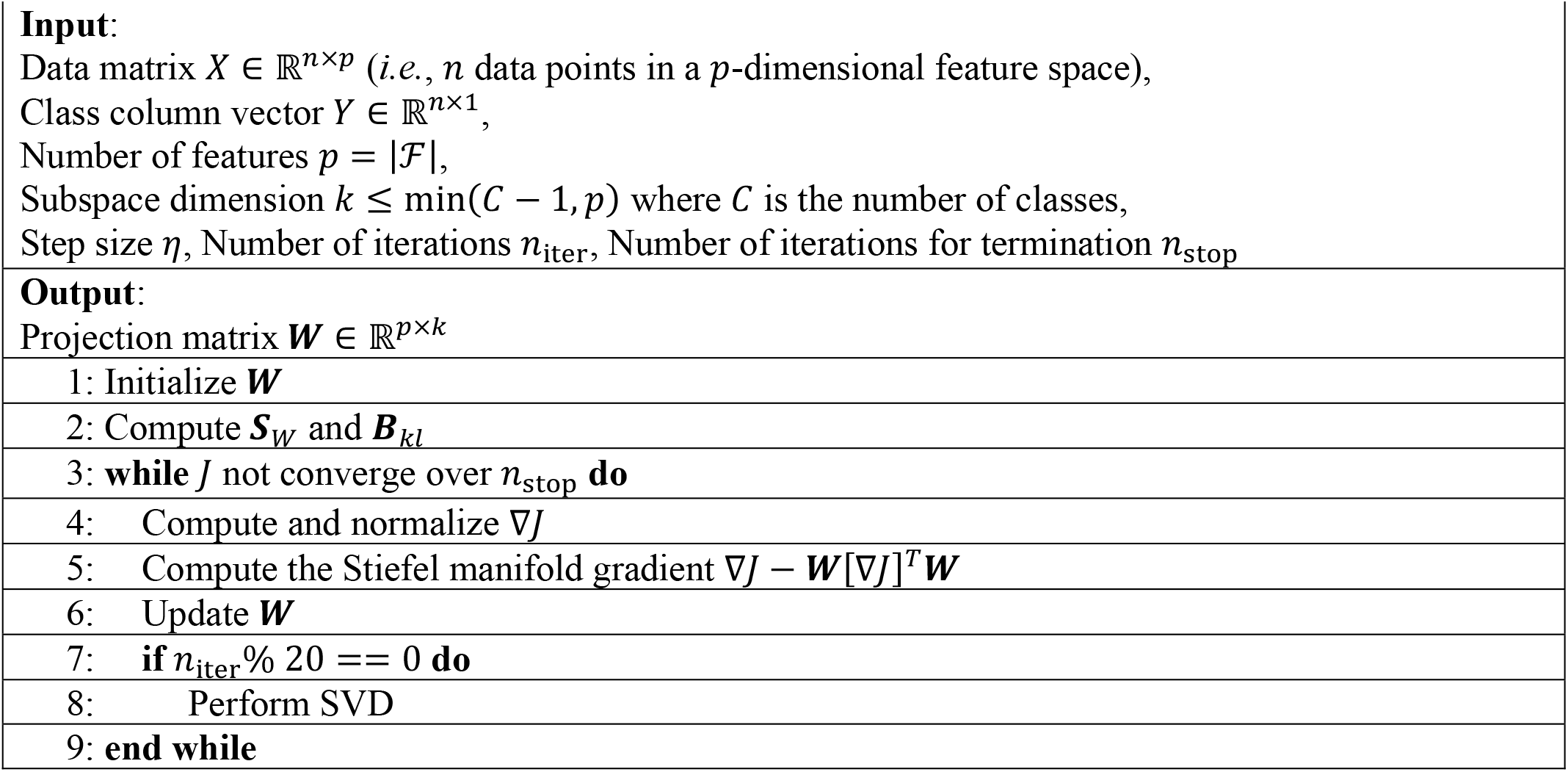

### Measures of Class Separation

To evaluate the distinguishability of CVs, we employed two metrics: the Gini Impurity Score (GIS), which relies on local information (i.e., neighbors of each data point) to examine overlaps between data distributions, and the Distance-based Separability Index (DSI)^71^, which provides global information by assessing both the overall data distributions and the distributions of pairwise distances between data points. To implement the GIS, we overlayed a rectangular grid onto a 2D space of CVs (obtained from one of the above DR analyses). The grid comprised 10,000 equally proportioned boxes, each measuring 0.4 by 0.4 units. Subsequently, we calculated the Gini Impurity^72-73^ *G* for each box using the formula:

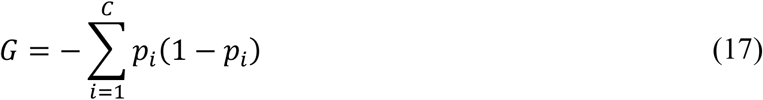

Here, *C* represents the total number of classes (in this study – three), and *p*_*i*_ is the fraction of data points within a given grid box that are ascribed to class *i*. The GIS metric then was defined as the percentage of the grid boxes with non-zero *G* values. A high GIS value is, thus, indicative of a substantial class overlap and, consequently, poor class separation.

The DSI^71^ for each class was computed as:

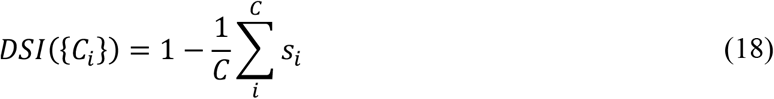

where *C*_*i*_ and *C* represent class *i* and the number of classes, respectively, and *s*_*i*_ is the Kolmogorov-Smirnov (KS) distance for class *i* defined as:

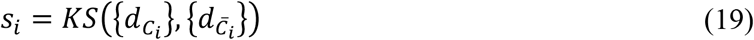

Here, 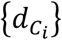 and 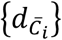 represent the intra-class (ICD) and between-class (BCD) distance sets for class *i*, respectively. The DSI serves as a normalized indicator of data separability (i.e., *DSI* ∈ (0,1)). Similar to GIS, a large DSI value suggests a substantial class overlap and therefore poor class separability.

### Deep Convolutional Neural Network (DCNN) Algorithm

We implemented a non-linear strategy using a DCNN classification algorithm for pattern recognition analysis of MD trajectories, detailed in Ref 36 (the python code is freely available at https://github.com/weinsteinlab/DNN) and utilized in our prior work on MFSD2A^30^. Here we used the DCNN method to classify three distinct metastable states of MFSD2A (OFS, OcS, and IFS). The protocol involves converting three-dimensional MD trajectories into a two-dimensional image format by mapping the x, y, and z coordinates of each atom to the red, green, and blue (RGB) values of pixels, respectively. This transformation allows for the application of image-based deep learning classification algorithms to structural biological data without compromising the integrity of the information. The input for the DCNN was ∼18 μs long MD trajectory frames (500 ns per 12 independent replicas for each state, i.e., ∼6 μs per state). The data was randomly split into a sample amount at a 56:24:20 ratio and outputted every 0.8ns, resulting in 12,701 training, 5,445 validation, and 4,536 test samples. The DCNN achieved 98.4% accuracy on the training set (95.5% for OFS, 99.9% for OcS and 99.8% for IFS), 97.9% on the validation set (95.1% for OFS, 99.6% for OcS and 99.3% for IFS), and 97.7% on the test set (93.9% for OFS, 99.7% for OcS and 99.5% for IFS). To avoid bias from protein orientation, input frames underwent positional and orientational scrambling, after which the coordinates of the TM segments (see above) were extracted and saved into separate trajectory files. The spatial coordinates were converted into a visual format for the DCNN, mapping *x, y, z* coordinates to RGB values in images sized (72,72,3) for the first 5184 TM atoms. The DCNN architecture included a convolutional layer with 2-stride, and a densely connected convolutional network (DenseNet)^74^ with three blocks of 6, 12, and 24 layers, starting with 96 filters and a growth rate of 48 per layer, halved in each subsequent block. The network was implemented using Keras^75^ with TensorFlow^76^, following an established architecture^77^. Training involved 30 epochs, with a dropout rate of 0.2 and weight decay of 0.00002. Sensitivity analysis employed gradient computation via the Keras-vis^78^ visual saliency package with guided backpropagation. Feature importance was evaluated using the Fisher-Jenks algorithm^79^ (https://github.com/mthh/jenkspy) to delineate saliency boundaries for the gradients of the OFS, OcS and IFS classes. Thus, for each of the classes (OFS, OcS, and IFS) using Fisher-Jenks algorithm a threshold at the upper tercile was established to discern the most salient features (see Results). The procedure yielded the following saliency thresholds: 0.28 for OFS, 0.31 for OcS and 0.32 for IFS.

## RESULTS

### Labeling of MFSD2A Metastable States in MD Simulation Trajectories

In this study, our aim is to apply various DR approaches (see Methods) for extracting CVs from unbiased sampling of local fluctuations in the MFSD2A lysolipid transporter in three distinct functional states – OFS, OcS, and IFS. The purpose of these CVs is to enable automated structural classification of MFSD2A in these metastable states. The MFSD2A system was chosen here as a case study for its biological significance and intricate functional mechanism which has been well-characterized at the detailed molecular level^29-32^. To create conformational ensembles of MFSD2A in the three metastable states, we conducted 18 μs of unbiased atomistic MD simulations of MFSD2A in OFS, OcS, and IFS (each state simulated in 12 independent 500 ns-long replicas, i.e., for 6 μs cumulative time, see Methods for details).

As reported previously^30^, the three metastable states in MFSD2A can be broadly structurally distinguished by the following specific features (**Figure 2a-c**): 1) the arrangement of TM2/11 and TM5/8 pairs of helices which can be quantified by measuring the distance between C_*α*_ atoms of extracellular (EC) residues V200 (on TM5) and F329 (on TM8) and the distance between C_*α*_ atoms of EC residues S78 (on TM2) and T447 (on TM11), and 2) the arrangement of TM7 helix which can be conveniently described by measuring its separation on the extracellular (EC) side from TM1 helix (pairwise distance between F61 and A316 residues on TMs 1 and 7, respectively), and by assessing its positioning on the intracellular (IC) side with respect to TM11 helix (quantifying the dihedral angle of residue Y294 in TM7).

**Figure 2:**
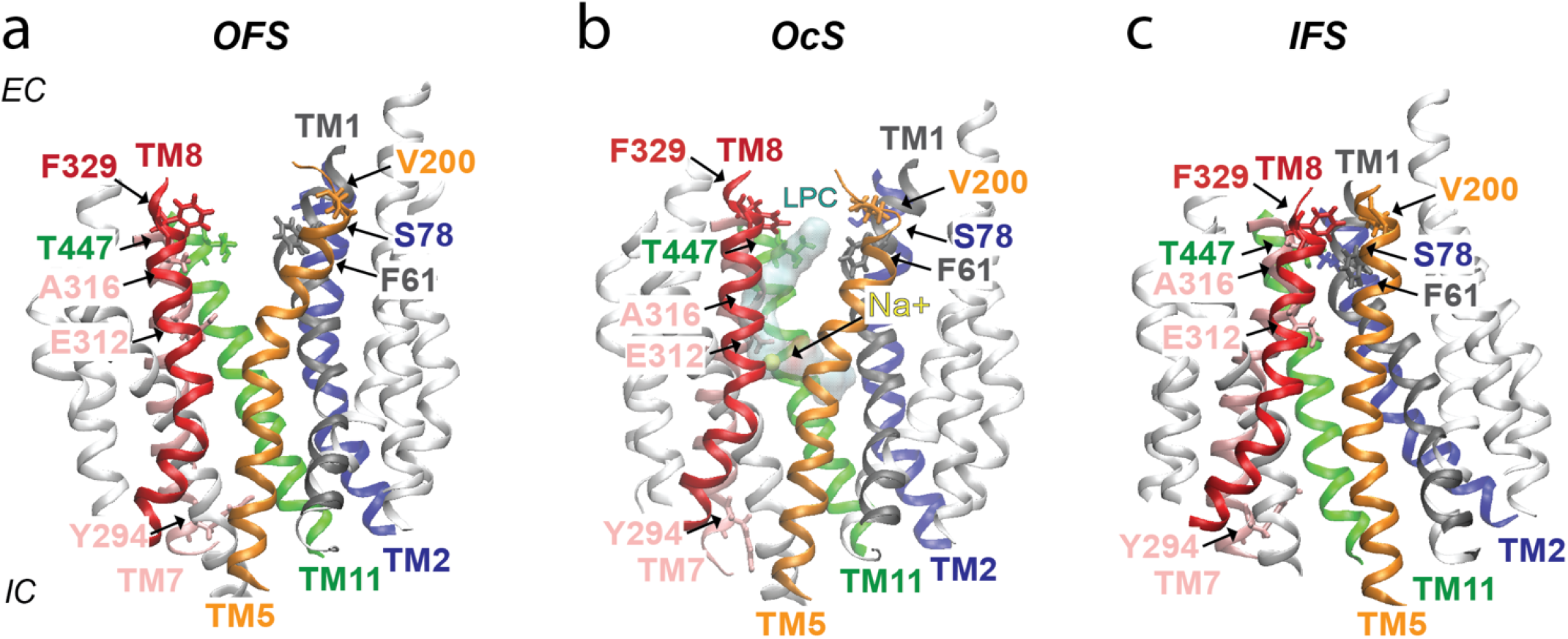
Side view (from the membrane plane) of ggMFSD2A in **(a)** OFS, **(b)** OcS, and **(c)** IFS. The OFS and IFS structures are the cryo-EM models (PDB IDs 7N98 and ligand-free 7MJS, respectively), while the OcS structure was predicted from our previous MD simulations^30^. TM2, 5, 8 and TM11 are shown in blue, orange, red, and green, respectively; TM1 and TM7 are shown in grey and pink, correspondingly. The remainder of the protein is colored in white. Residues F61, S78, V200, A316, F329, and T447 are colored according to the colors of the helices they belong and shown in licorice representation. The LPC-18:1 substrate in the OcS is shown in transparent surface representation. The Na^+^ ion, displayed as a yellow sphere, coordinates the central cavity residue E312 (in pink) in the OcS.

We used these measures to confirm that during our MD simulations, the transporter protein was dynamically sampled around the initial metastable states without transitioning between them. Indeed, as shown in **Figure 3b-e** and **Table 1**, the above structural parameters largely fluctuated around their respective values in the initial structural models. The overall stability of the protein structure in our simulations is further corroborated by the low values of RMSD of the backbone residues in TM helices across the trajectories. As shown in **Figure 3a**, the RMSD value for the OcS predominantly falls below 1 Å, while for the IFS and OFS simulations, the peak RMSD values are less than 1.5 Å relative to their respective initial conformations. Collectively, these analyses confirm the structural stability of MFSD2A in the simulations and allow us to unambiguously label the MD trajectory frames according to OFS, OcS, and IFS classes.

**Table 1:** Quantitative Analysis of MFSD2A MD Simulations Using Established Conformational State Metrics: The table summarizes the mean values for the RMSD of the protein TM helical bundle backbone atoms (row 1), for the key interatomic distances (rows 2-4), and for the Y294 dihedral angle (row 5) obtained from MD simulations of MFSD2A in the three metastable states (OFS, OcS, and IFS, see also **Figure 3**). These mean values are compared to the respective quantities measured in the initial structural models.

| Metric |  | Selection | OFS<br>MD<br>Simulations | *<u>OFS</u>*<br>*<u>Initial</u>*<br>*<u>Structure</u>* | OcS<br>MD<br>Simulations | *<u>OcS</u>*<br>*<u>Initial</u>*<br>*<u>Structure</u>* | IFS<br>MD<br>Simulations | *<u>IFS</u>*<br>*<u>Initial</u>*<br>*<u>Structure</u>* |
| --- | --- | --- | --- | --- | --- | --- | --- | --- |
| 1 | RMSD (Å) | TM helices | Mean: 1.51 | — | Mean: 0.93 | — | Mean: 1.50 | — |
| 2 | V200-F329<br>distance (Å) | Cα-Cα<br>atoms | Mean: 18.71 | **20.16** | Mean: 14.84 | **14.20** | Mean: 10.79 | **9.42** |
| 3 | S78-T447<br>distance (Å) | Cα-Cα<br>atoms | Mean: 12.97 | **12.19** | Mean: 10.56 | **11.16** | Mean: 5.78 | **5.45** |
| 4 | F61-A316<br>distance (Å) | Cα-Cα<br>atoms | Mean: 15.24 | **15.74** | Mean: 13.96 | **14.13** | Mean: 7.85 | **7.59** |
| 5 | Y294<br>dihedral<br>angle (°) | C-Cα-Cβ-Cγ<br>atoms | Mean: 56.49 | **53.23** | Mean: 172.38 | **179.42** | Mean: 47.44 | **39.47** |

**Figure 3:**
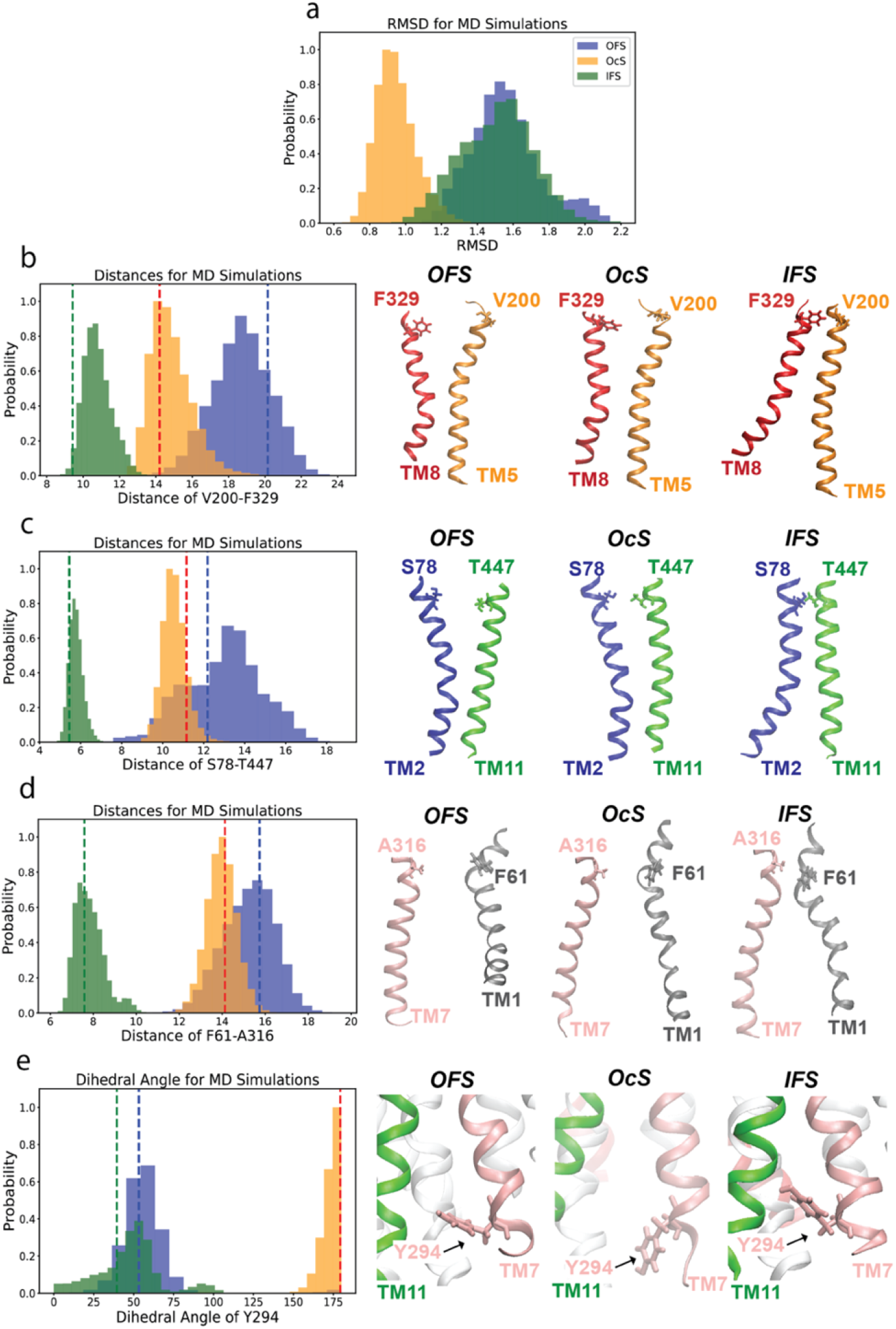
Analysis of MFSD2A Metastable States in MD Simulations. **(a)** Distribution of RMSD values for the backbone atoms of MFSD2A’s TM helices in the MD trajectories. The data for OFS (blue), OcS (orange), and IFS (green) are shown in separate histograms. **(b-d)** Probability distributions of V200-F329 (b), S78-T447 (c), and F61-A316 (d) distances in the MD trajectories. The data for OFS, OcS, and IFS are shown in separate histograms as in panel a. **(e)** Probability distributions of the dihedral angle of residue Y294 in the three metastable states (shown in separate histograms as in panel a). The vertical dashed lines in panels b-e represent the values of the measured quantities in the initial frame of the respective MD trajectory. The structural snapshots accompanying panels b-d illustrate the arrangement of TM5-TM8, TM2-TM11, and TM1-TM7 helical pairs, respectively, in OFS, OcS, and IFS, with the key residues shown as sticks and labelled. The snapshots accompanying panel e show the positioning of residue Y294 in the three metastable states, with TM11 and TM7 helices depicted in green and pink colors, respectively.

### Data Filtering and Feature Selection

After each trajectory frame was assigned to one of the three classes (OFS, OcS, or IFS), in each frame we calculated pairwise distances between the TM bundle residues using either geometric centers of the residues, 𝒟_RE_, or the *C*_*α*_ atoms, 𝒟_CA_ (see Methods and **Figure 1**). Next, using the linear DR algorithms described in Methods (see also below), we sought to combine these distances into CVs that would allow classification of the MD data according to the three metastable states. Prior to the DR step, we performed feature selection procedure (**Figure 1**), during which 𝒟_RE_ and 𝒟_CA_ datasets were trimmed down to a smaller number of features to ensure that only the most relevant features from these large datasets are passed on to the DR algorithms. By doing so, the CVs obtained at the DR stage will be dependent only on a small number of pertinent distance variables, and thus will be expected to be easily interpretable in the detailed structural context. As described in Methods, we used several alternative approaches for the feature selection procedure in order to ensure the robustness of the final set of chosen features. These included filter-based methods, such as the χ^2^ test and Fisher scores combined with the AMINO protocol, and wrapper methods, such as BPSO-SVM and MPSO-SVM.

For all the above methods, we first identified and removed constant and quasi-constant features by calculating the variance of each feature and eliminating those whose variance was below a predetermined threshold using the Kneedle algorithm (see Methods, **Figure S1**). This resulted in trimming 𝒟_RE_ and 𝒟_CA_ datasets to 12,371 and 16,639 number of features, respectively. Following this variance-based filtering, the χ^2^ or Fisher scores were calculated to the remaining features followed by the AMINO algorithm.

#### 1. Features identified with χ^2^*Test* - AMINO *protocol*

We calculated the χ^2^ scores as described in Methods, using a score threshold of 16,910 for the 𝒟_RE_ distance set and 17,227 for the 𝒟_CA_ set (**Figure S1c,d**). This procedure resulted in the reduction of the 𝒟_RE_ and 𝒟_CA_ datasets to 1460 and 1594 features, respectively. Following the χ^2^ test, we employed the AMINO algorithm to further refine our feature set, by eliminating the features that share high mutual information (see Methods), i.e., are redundant features. This yielded 5 molecular features for the 𝒟_RE_ set, and 4 – for the 𝒟_CA_ set.

#### 2. Features identified with Fisher Score *Test* - AMINO *protocol*

The application of the Fisher score analysis to the variance-based trimmed dataset (see above) resulted in identification of the top 4014 features for 𝒟_RE_ and 3093 features for 𝒟_CA_. As described in Methods, for this procedure we used a score threshold of 5.87 for the 𝒟_RE_ distance set and 6.47 for the 𝒟_CA_ set (**Figure S1e,f**). Similar to the χ^2^ test, we applied the AMINO algorithm to the dataset generated by the Fisher score analysis, to further refine our feature set. The procedure identified 5 molecular features for the 𝒟_RE_ set, and 8 – for the 𝒟_CA_ set.

#### 3. Features identified with BPSO-SVM method

The iterative application of BPSO-SVM algorithm to the variance-based trimmed dataset described above, identified 5 key molecular features for both 𝒟_RE_ set 𝒟_CA_ sets.

#### 4. Features identified with MPSO-SVM method

As described in Methods, the MPSO analysis was performed on the variance-based trimmed dataset under conditions of strong (MPSO-S) or loose (MPSO-L) regularization. With MPSO-S, 5 features were identified for the 𝒟_RE_ dataset and 4 for the 𝒟_CA_ dataset, while the MPSO-L algorithm found 16 features for the 𝒟_RE_ dataset and 11 for the 𝒟_CA_ dataset.

**Table 2** summarizes the data from all the above feature selection methods. The identified features include pairwise distances that reflect on differential positioning of specific TM domains in OFS, OcS, and IFS as previously reported in the literature. Indeed, as presented earlier^29,30^ and described above (see **Figure 1**), OFS→OcS→IFS transitions are accompanied by conformational changes in TM2/TM11 and TM5/TM8 pairs of helices such that these helical pairs assume V-shape in OFS, inverted V-shape in IFS, and relatively parallel orientation in OcS.

**Table 2:** Summary of identified features derived from each feature selection methods. The table lists selected features (pairwise distances) for MFSD2A 𝒟_RE_ and 𝒟_CA_ datasets determined by filter methods (χ^2^-AMINO and Fisher-AMINO) and wrapper method (BPSO-SVM and MPSO-SVM with strong and loose regularization). Residues located in TM2, 5, 7, 8 and TM11 are shown in blue, orange, pink, red, and green, respectively.

|  | *Filter Method* |  | *Wrapper Method* |  |  |
| --- | --- | --- | --- | --- | --- |
| | $\chi^2$-AMINO | Fisher-AMINO | BPSO | MPSO | |
|  |  |  |  | MPSO-S | MPSO-L |
| *Res-Res Distances ($\mathcal{D}_{RE}$)* | V139.I470<br>**I80**.I468<br>**V200**.A388<br>L110.**Y294**<br>**V83**.I386 | **F82**.**Y294**<br>G137.**I199**<br>P159.I410<br>T238.L470<br>**I80**.L470 | **F75**.**Y294**<br>**F75**.**S305**<br>**G96**.**F427**<br>**Y294**.A391<br>**E424**.L491 | G47.**I295**<br>**L81**.**F308**<br>G137.G250<br>**W87**.**Y294**<br>L110.**Y294** | **G84**.A390; **W87**.**V441**<br>I123.V479; V46.**I426**<br>G137.D237;<br>**Y180**.**F350**<br>L110.K490;<br>Q138.A241<br>**E173**.Y361;<br>**V95**.**K296**<br>Q138.**G444**;<br>F149.**Y294**<br>**Y294**.L333; **R85**.F350<br>**R181**.L415;<br>A246.**E328** |
| *CA-CA Distances ($\mathcal{D}_{CA}$)* | **V200**.A388<br>**I80**.I468<br>W113.**Y294**<br>**L81**.G392 | **F75**.**A307**<br>G137.**Q198**<br>G137.**I199**<br>T238.L470<br>V139.I468<br>L66.**L310**<br>F145.L333<br>**I199**.**K202** | C39.**Y294**<br>**E173**.K414<br>**V200**.A367<br>A257.S404<br>**G292**.E465 | G137.**G192**<br>I115.**A293**<br>P112.**S429**<br>S234.K471 | I114.**L334**;<br>G137.L235<br>**Q171**.**I344**;<br>**V200**.**T304**<br>W141.V474;<br>F61.**T447**<br>D68.**L310**; I123.**L310**<br>C43.**S429**; **F97**.**Y294**<br>A246.**A293** |

Our analysis revealed that 17 out of 68 features (∼25%) reflected on TM2 positioning (highlighted in **Table 2** in blue), and 8 out of 68 features (∼12%) described TM11 positioning (highlighted in **Table 2** in green). Notably, the features G96.F427 and W87.V441 indicate distance changes specifically between TM2 and TM11. Furthermore, 14 of the 68 features (∼21%) describe TM5 location (denoted in orange in **Table 2**), and 6 of the 68 features (∼9%) reflected on TM8’s positioning (highlighted in **Table 2** in red), with the residue pairs Y180.F350 and Q171.I344 quantifying relative positioning of TM5 and TM8 helices.

Another important structural discriminant between OFS, OcS, and IFS is positioning of TM7^30^, as described above (**Figure 3e**). Consistent with this notion, 23 of the 68 features (∼34%) in **Table 2** relate to TM7 arrangement (marked in pink in **Table 2**). Furthermore, within these 23 features, 11 include residue Y294 on TM7. Conformational changes in this residue were identified to play an important mechanistic role during OFS→OcS→IFS^30^ (**Figure 3e**).

Interestingly, **Table 2** also contains structural features that have not been highlighted in the previous studies and include mostly outer helices of the MFSD2A protein bundle, such as TM4, TM6 and TM12. Thus, our analysis identifies as important features for structural discrimination of the three functional states distances between TM4-TM6 (G137.G250, G137.D237, Q138.A241, G137.L235), TM6-TM12 (T238.L470, T238.L470, A257.S404, S234.K471), TM4-TM12 (V139.I470, V139.I468, W141.V474), TM3-TM12 (I123.V479, L110.K490), TM4-TM8 (F145.L333) and between TM4-TM10 (P159.I410).

Taken together, our results show that the majority of the features identified by the different feature selection approaches are consistent with the structural properties that were shown previously to be modulated by state-to-state transitions in MFSD2A. Identification of these features from our feature selection algorithms in an automated manner affirms the robustness of the methods for their ability to discriminate between OFS, OcS, and IFS. To establish the most significant structural motifs for classification and to construct CVs from the sets of features in **Table 2**, we next applied various linear DR techniques to the features.

### LDA-based DR methods combined with MPSO feature selection algorithms result in the best classification performance

#### a. CA-CA Distances (𝒟_CA_)

Figure 4. illustrates the projections of the 𝒟_CA_ data points from the OFS, OcS, and IFS trajectory frames onto the subspaces defined by the first two CVs derived from the array of LDA-based DR algorithms as well as from PCA. The results show that the first CV (LD1 or PC1) generally separates IFS from the other two states (OcS/OFS), whereas the second CV (LD2 or PC2) mostly separates OcS from OFS (see also below). Overall, our results show that different combinations of DR and feature selection methods yield varying levels of class separation in the subspace. To quantify the degree of class separability, we employed two metrics described in Methods: the GIS (providing more local information focusing on overlaps between data distributions) and the DSI (providing more global information focusing on overall data distributions).

Analysis of the GIS for the 𝒟_CA_ dataset, shown in the upper panel of **Figure 5**, reveals several trends. First, we observed that PCA tends to exhibit a greater degree of class overlap in comparison to the LDA-based methods, irrespective of the feature selection technique employed. Second, the LDA-based algorithms demonstrate a relatively consistent pattern in their ability to distinguish classes; BPSO and MPSO methods are associated with lower GIS values compared to the χ^2^ test and Fisher scores. Additionally, features identified through the χ^2^ test consistently result in the highest degree of class overlap, regardless of the DR algorithm applied. Importantly, the second CV (PC2 or LD2 in **Figure 5**) contributes minimally to class separation when the χ^2^ test is used. In contrast, features selected by MPSO are distinguished by the least extent of class overlap, suggesting higher class separability. Furthermore, MPSO-L is seen to perform marginally better compared to the MPSO-S. This should not be surprising, as each CV from MPSO-L incorporates larger number of features (11) compared to the CVs from the MPSO-S (only 4 features).

**Figure 4:**
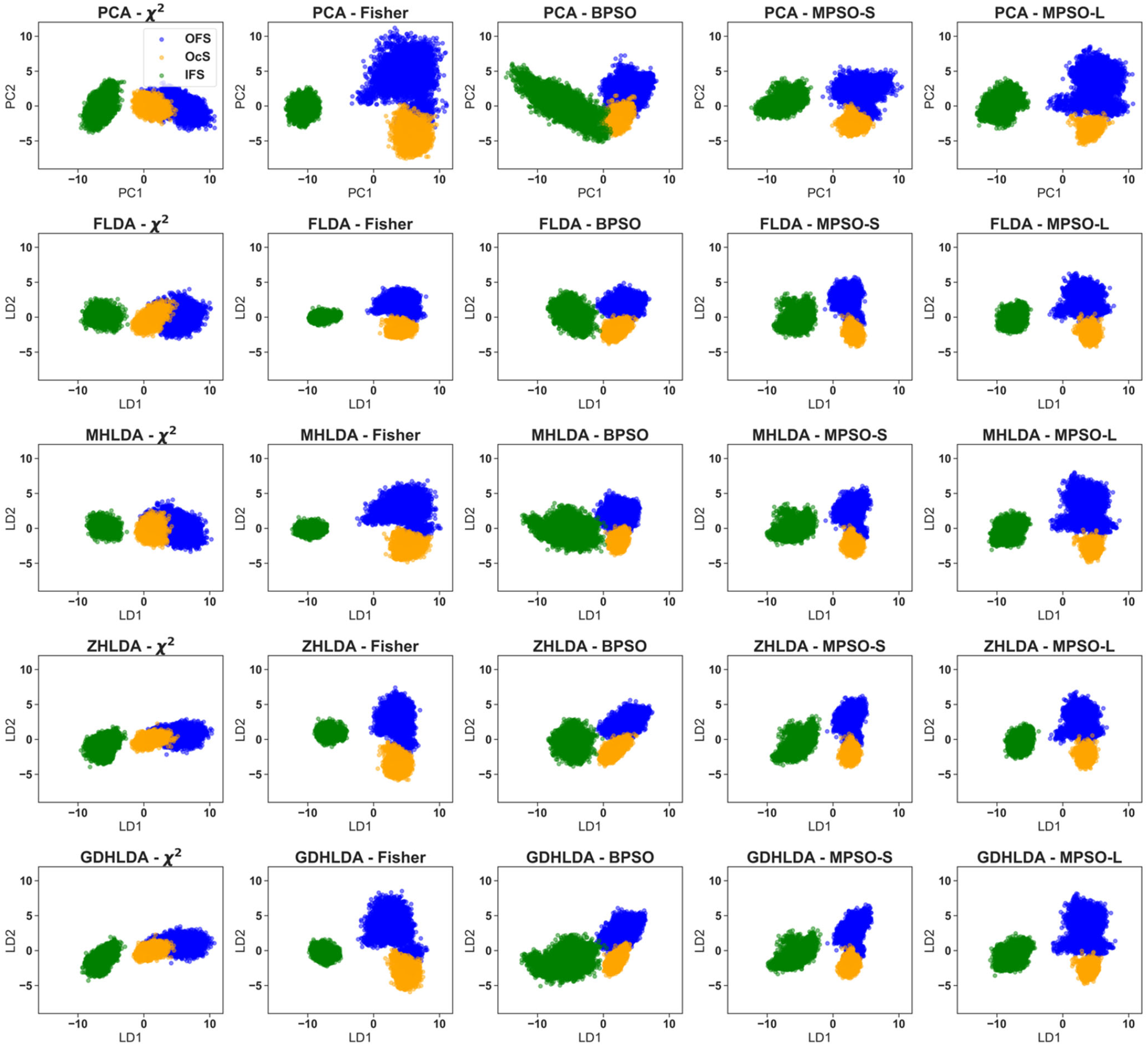
DR results for the *𝒟*_CA_-Dataset using various feature selection methods. Each subplot represents a two-dimensional scatter plot of the trajectory frames projected onto the first two principal components (LD or PC) for different combinations of feature selection methods and DR techniques. Each column represents a different feature selection method: χ^2^, Fisher, BPSO, MPSO-S or MPSO-L. Each row represents different DR techniques: PCA, FLDA, ZHLDA, MHLDA, and GDHLDA. To facilitate visual assessment of class separability achieved by each method, the data points are color-coded as follows: blue for OFS, orange for OcS, and green for IFS.

**Figure 5.**
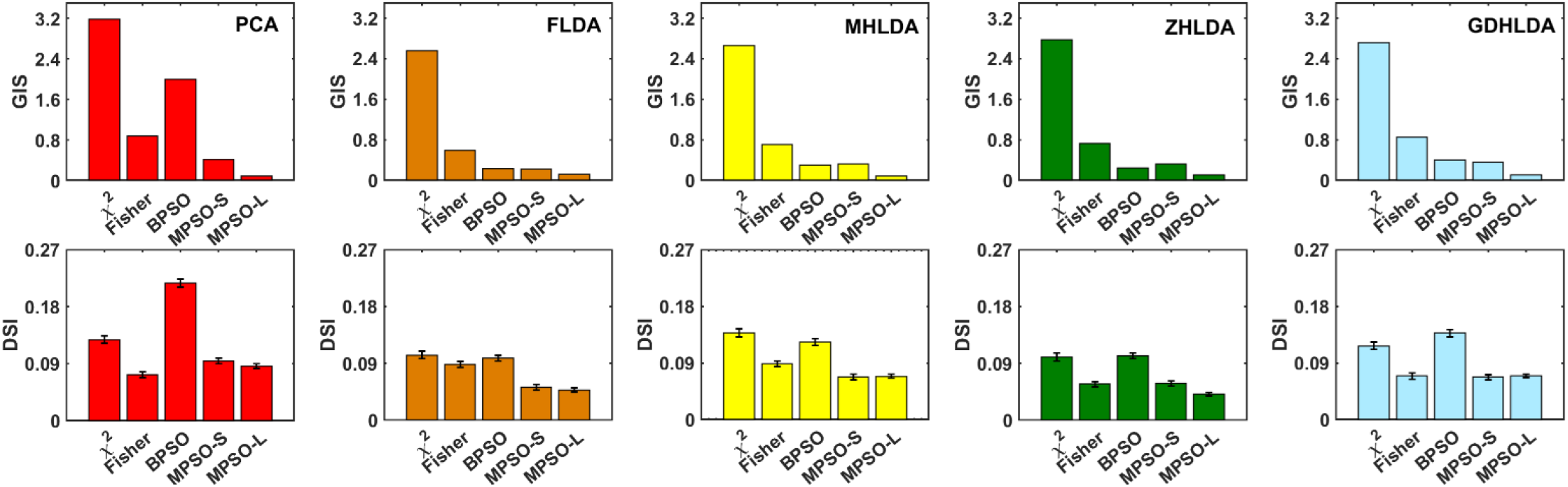
Plots of GIS and DSI values for the *𝒟*_*CA*_ dataset. The upper panel shows GIS values, as the proportion of the grid covered by impure boxes, and the lower panel shows DSI values for class separation achieved by features derived from χ^2^, Fisher, BPSO, MPSO-S, and MPSO-L. The bar graphs are colored in red for PCA, brown for FLDA, yellow for MHLDA, green for ZHLDA, and light blue for GDHLDA. The error bars in the lower panel were estimated using bootstrapping with 100 iterations and a subsample size of 1000.

Results of the DSI scores for the 𝒟_CA_ dataset, depicted in the lower panel of **Figure 5**, yield overall similar insights as the GIS data. Thus, the DSI values for PCA are higher compared to the LDA-based methods, suggesting poorer class separation. Second, the plots corresponding to the LDA-based methods exhibit a highly consistent pattern depending on the chosen feature selection technique. Specifically, the χ^2^ test and BPSO feature selection methods yield lower class separability across all DR algorithms whereas the MPSO performed the best. Taken together, the GIS and DSI metrics suggest that the LDA-based methods combined with the MPSO-based feature selection algorithms are the most optimal for the 𝒟_CA_ data classification.

#### b. RES-RES Distances (𝒟_RE_)

Results from the similar analyses performed on the 𝒟_RE_ dataset are shown in **Figures 6-7**. The projections of the 𝒟_RE_ data points onto the subspaces defined by the first two CVs derived from PCA and LDA-based methods (**Figure 7**) reveals that the class separation achieved for the 𝒟_RE_ dataset is overall superior to that of the 𝒟_CA_ dataset. The GIS results for the 𝒟_RE_ dataset (displayed in the upper panel of **Figure 7**) show the patterns akin to those in the 𝒟_CA_ dataset, with PCA tending to overall manifest higher class overlap relative to the LDA-based methods (with the exception of the Fisher score-based feature selection method). In addition, FLDA and ZHLDA, which do not use the harmonic mean in calculating the between-class scatter matrix, exhibit parallel trends. Conversely, MHLDA and GDHLDA, which do employ the harmonic mean for the between-class scatter matrix, display similar patterns. As with the 𝒟_CA_ dataset, the χ^2^ test for feature selection for the 𝒟_RE_ dataset results in the worst class separation across all DR algorithms, while the MPSO-based methods yield the best classification performance.

**Figure 6:**
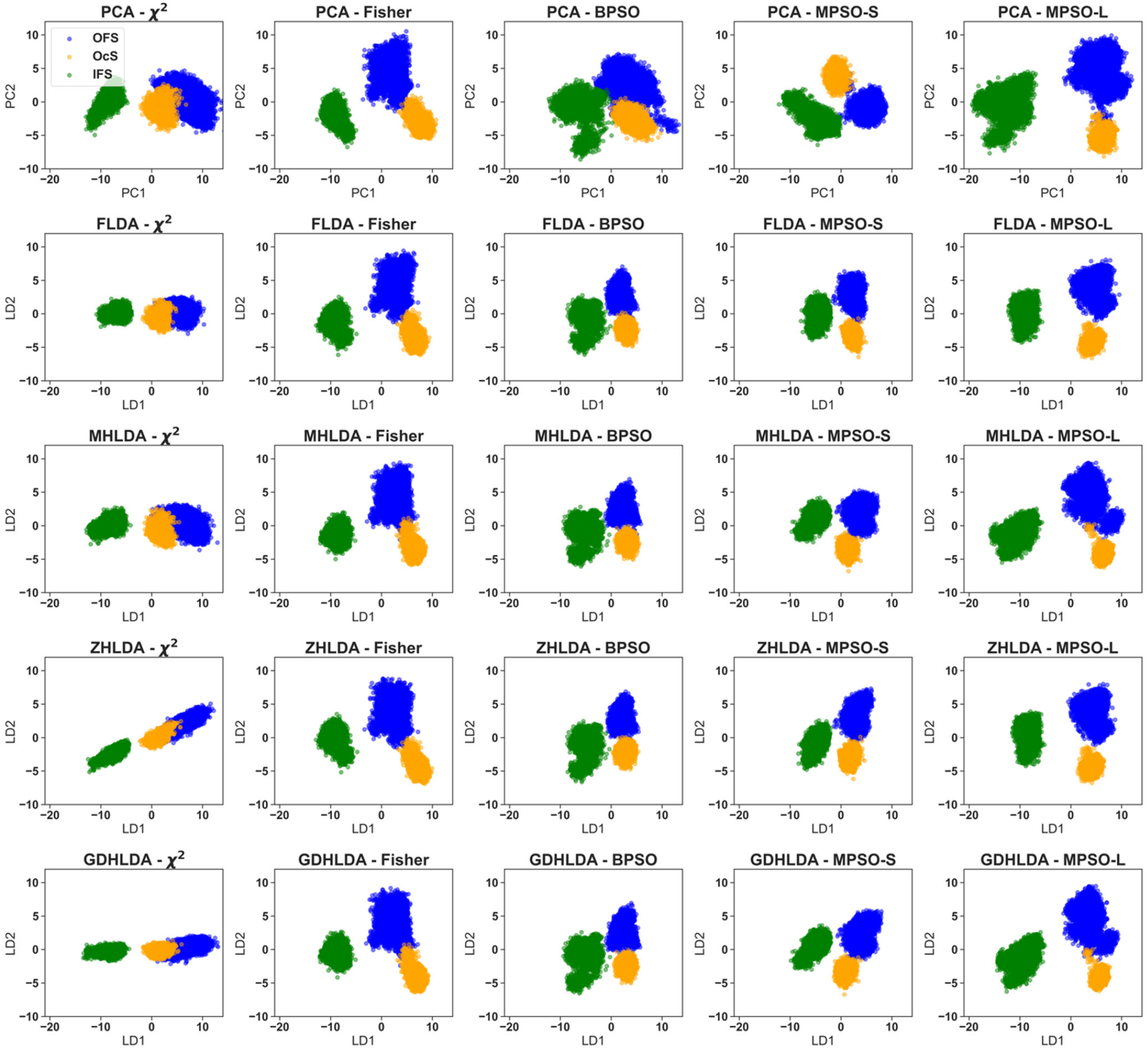
DR results for the *𝒟*_RE_-Dataset using various feature selection methods. Each subplot represents a two-dimensional scatter plot of the trajectory frames projected onto the first two principal components (LD or PC) for different combinations of feature selection methods and DR techniques. Each column represents a different feature selection method: χ^2^, Fisher, BPSO, MPSO-S or MPSO-L. Each row represents different DR techniques: PCA, FLDA, ZHLDA, MHLDA, and GDHLDA. To facilitate visual assessment of class separability achieved by each method, the data points are color-coded as follows: blue for OFS, orange for OcS, and green for IFS.

**Figure 7.**
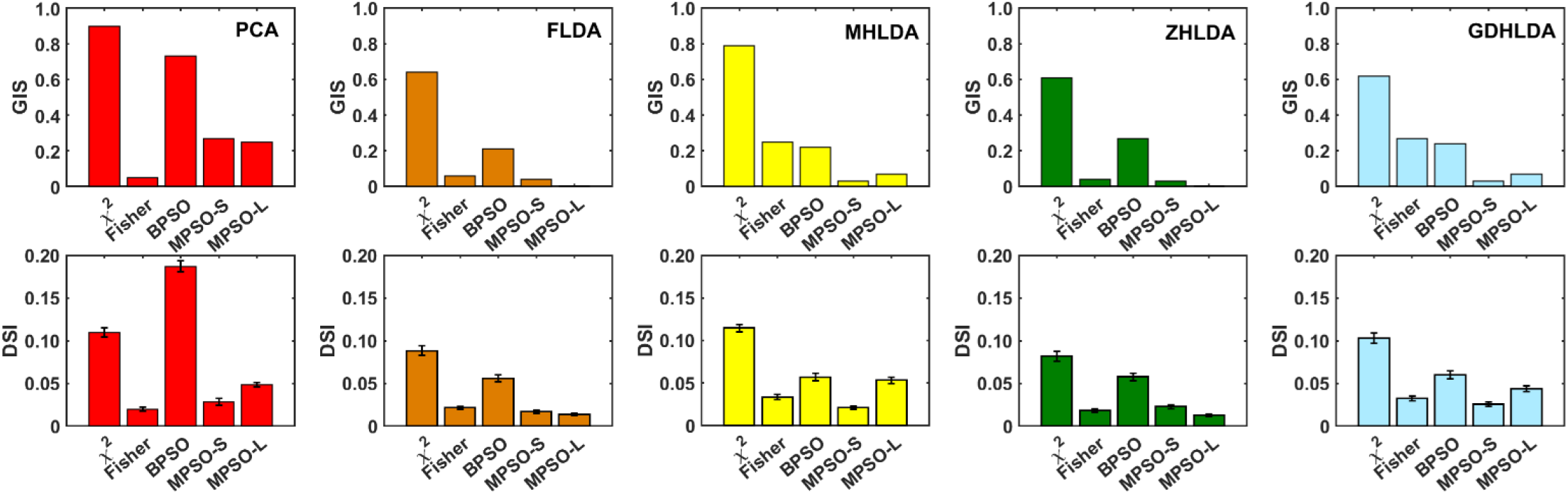
Plots of GIS and DSI values for the *𝒟*_*RE*_ dataset. The upper panel shows GIS values, as the proportion of the grid covered by impure boxes, and the lower panel shows DSI values for class separation achieved by features derived from χ^2^, Fisher, BPSO, MPSO-S, and MPSO-L. The coloring scheme is the same as in **Figure 5**. The error bars in the lower panel were estimated using bootstrapping with 100 iterations and a subsample size of 1000.

Taken together with the DSI values for the 𝒟_RE_ dataset (lower panel of **Figure 7**), our results reveal that the 𝒟_RE_ dataset is characterized by better class separation compared to the 𝒟_CA_ dataset. Furthermore, for both 𝒟_RE_ and 𝒟_CA_ datasets the LDA-based methods generally outperform PCA in the classification task, and the MPSO-based feature selection algorithms show overall best classification performance. We also note that GDHLDA generally achieves a level of class separability that is comparable to the other LDA methods.

### Comparative Analysis of Key Features Contributing to MFSD2A CV Design

We next analyzed contributions of the individual features to CV1 and CV2. **Figure S3 and S4** show the weights of the features and their importance scores for CV1 and CV2 for each DR approach for the 𝒟_CA_ and 𝒟_RE_ datasets, respectively. The features identified as having the most significant contributions to the CVs (i.e., those with the highest weights totaling more than 0.5) are listed in **Table S1-S4** and summarized in **Table 3**. These features show a consistent pattern in their structural localization. Indeed, as can be seen from **Table 3**, ∼73% of the highest contributing features are associated with TMs 2, 5, 7, and 11, the helical segments whose positioning was previously found to change prominently during the state-to-state transitions in ggMFSD2A. Specifically, 18 such features (∼33% of the highest contributors) were associated with TM7, 12 (∼22%) with TM5, 11 (∼20%) with TM2, and 6 (∼11%) with TM11, suggesting that the distance changes involving the residues from these TMs strongly contribute to CV1 and CV2. Interestingly, **Table 3** reveals that significant portion of the highest contributing features contains residues that reside on other TMs as well, such as TM4 (16.3%) and TM6 (12.7%), suggesting that the structural differences between the three functional states extend globally to the TM bundle.

**Table 3:**
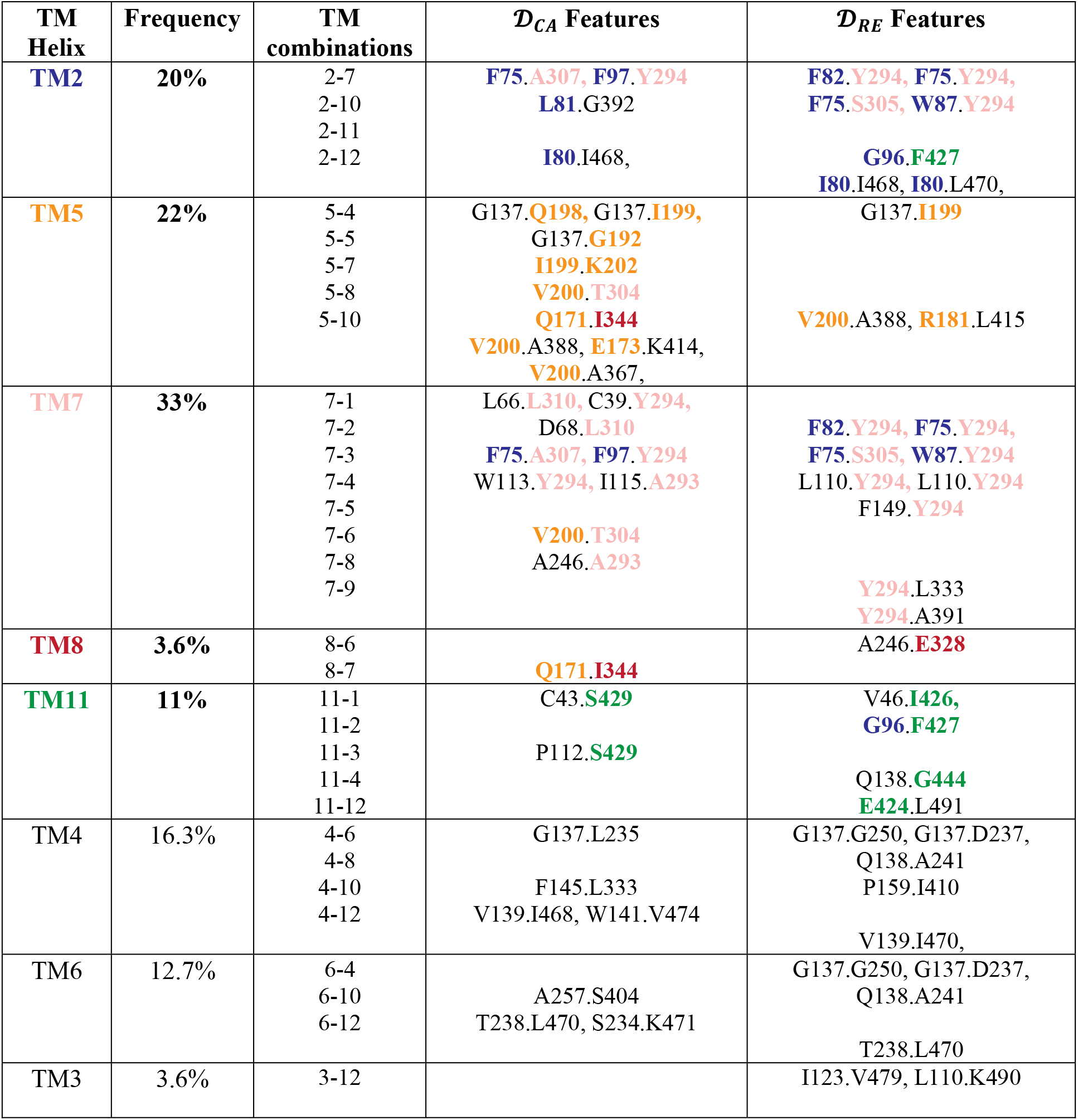
The most significant features in CV1/CV2 obtained from the DR analysis of MD data. The table enumerates the most significant features for the 𝒟_RE_ and 𝒟_CA_ datasets, specifying their association with TM helices. The frequency column denotes the percentage of features contained in each TM helix. TMs 2, 5, 7, 8, and 11 are color-coded in blue, orange, pink, red, and green, respectively.

Further analysis of the 𝒟_CA_ dataset reveals specific interhelical interactions that contribute the most to the individual CVs. For CV1, which separates IFS from OcS/OFS (**Figure 4**), we observed contributions from multiple helical pairs (TMs 1-5 paired with TMs 7-10 and 11), consistent with the global structural rearrangements the protein undergoes entering the IFS. Still, we find TM1-TM7 interactions being particularly frequent (14%) in CV1. Conversely, for CV2, which predominantly separates OcS from OFS (**Figure 4**), the features derive from a relatively smaller set of interhelical interactions (involving mostly TMs 4, 5, 6, 7, and 10), consistent with more subtle conformational differences between these two states^30^. The most common features in CV2 represent distances between TM4-TM5 helical pairs. Certain interhelical interactions are prominently featured in both CV1 and CV2. These include distances between TM1-TM7, TM5-TM10, TM6-TM10, and TM6-TM7 suggesting that relative positioning of these helical pairs vary significantly between all the three states.

Comparative analysis between the 𝒟_CA_ and 𝒟_RE_ datasets enables us to identify how differently/similarly collective repositioning of the helical segments and more localized side-chain level structural rearrangements contribute to the classification of the functional states. To this end, it is notable that both datasets showed a similar proportion of TM7 features (32% for 𝒟_CA_ and 33% for 𝒟_RE_), with a notable difference in capturing Y294 (89% of TM7 features in 𝒟_RE_ versus 33% of TM7 features in 𝒟_CA_), indicating that positioning of Y294 side-chain is an important factor for the classification, consistent with our previous findings^30^.

Interestingly, some of the highest contributing features in **Table 3** are reflective of local secondary structure differences between the OFS, OcS, and IFS. Thus, our analysis found that the EC ends of TM4 and TM5 helices (residue segments 137-138 and 198-202, respectively) adopt distinct secondary structures in the three states (**Figure S6**). These local structural changes likely influence our predictions (and some redundancies, such as the presence of G137.Q198 and G137.I199 distances), resulting in a high ranking of the features containing these residue segments.

### DCNN Reveals TM6 and TM7 as Crucial for Classifying MFSD2A Metastable States

To better evaluate the performance of the above linear classification models, we compared their predictions to those obtained from a non-linear approach using the supervised DCNN algorithm for pattern recognition analysis of MD trajectories (see Methods)^36^. This DCNN method had previously demonstrated high accuracy in classifying the OFS and OcS states of MFSD2A identifying TM7 as a key molecular feature for the classification^30^. Expanding on these results, here we applied the DCNN to the MFSD2A trajectories in all three metastable states—OFS, OcS, and IFS (see Methods). The DCNN achieved 97.7% accuracy on the test set (93.9% for OFS, 99.7% for OcS and 99.5% for IFS). This high level of accuracy demonstrates the model’s robustness in classifying molecular conformations (**Figure 8a**). This is reinforced by the observed decrease in the loss function to value of 0.2 during the 30 epochs run (**Figure 8b**). To discern the key molecular features for DCNN classification, we employed a visual saliency sensitivity analysis^36^, **Figure 8c-e**.

**Figure 8.**
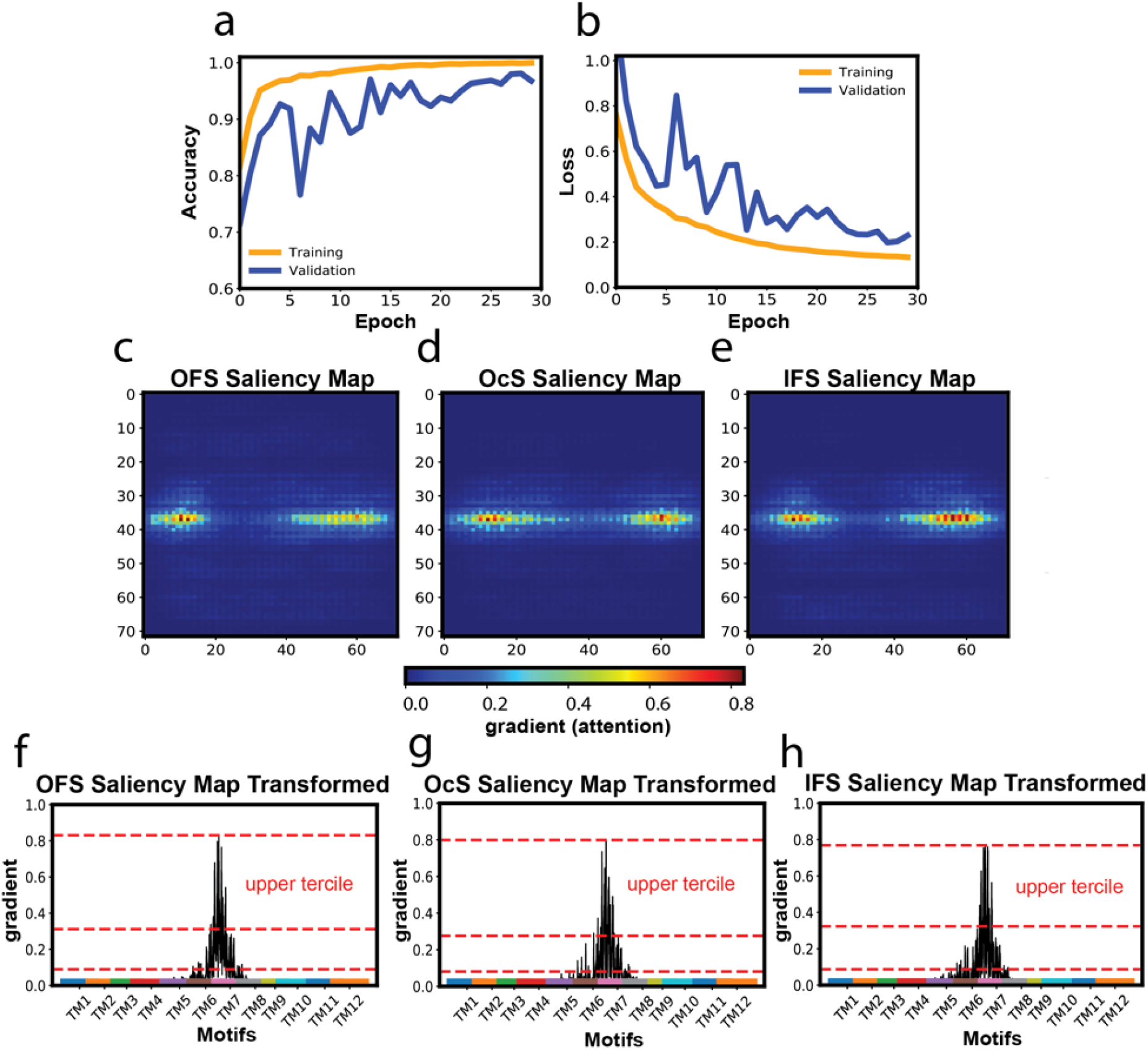
DCNN Model Performance and Saliency Mapping. **(a-b)** The accuracy (a) and loss (b) of the DCNN model plotted per epoch for both the validation and training sets. **(c-e)** Saliency (attention) maps generated by the DCNN for the OFS (c), OcS (d), and IFS (e). Each pixel corresponds to an atom in the TM domain of ggMFSD2A, organized by index from lowest (top left) to highest (bottom right). The color gradient indicates importance, with red highlighting atoms with the highest gradient and therefore greatest significance for classification. **(f-h)** The gradient values from the 2D maps in panels c-e plotted for all TM regions for OFS (f), OcS (g), and IFS (h), respectively. The most important regions of the protein for the classification were identified from the Fisher-Jenks clustering algorithm (see Methods). These regions, which fall within the upper tercile (indicated by the red dashed lines), include residues 249 to 263 on the IC end of TM6 and residue 292 to 306 on the IC end of TM7.

The saliency analysis identified residue segment 292-306 at the IC end of TM7 and residue stretch 249-263 at the IC end of TM6 (**Figure 8a-c** and **Figure S5**) as the most important features for classification. Interestingly, our analysis reveals consistency in the identified regions between the DCNN and the linear DR methods. Specifically, residues G250 and A257 in the TM6 segment, along with several residues in the TM7 region (**Table 3**), including Y294 residue, were identified by all the linear DR methods as well as by the DCNN. This consistency across different approaches suggests the importance of these structural motifs for the classification.

To confirm the significance of the features identified by the DCNN, the residue segments positions 249-263 and 292-306 were excluded from the input dataset, and the accuracy of the DCNN was reassessed. This exclusion resulted in a marked decrease in model accuracy, which dropped to 56.5% in the test set (68.5% for OFS, 39.5% for OcS, and 61.4% for IFS). Furthermore, a DCNN model that was exclusively trained on the limited dataset of the top salient residues (249-263 and 292-306) demonstrated exceptional performance, with an accuracy of 99.4% on the test set (99.7% for OFS, 99.6% for OcS and 98.9% for IFS). Taken together, these results confirm that the identified structural features are critical for accurate classification of MFSD2A metastable states.

### Interpretation of the CVs emerging from the DR analysis

We next sought to interpret the CVs identified from our DR analysis in more detailed structural context. To this end, we focused on the data obtained from the MPSO-S feature selection method since this approach resulted in the best overall separability of the classes with relatively small number of features, as described above (**Figure 5**). Thus, the identified features with this approach were G47.I295 (TM1/TM7), L81.F308 (TM2/TM7), G137.G250 (TM4/TM6), W87.Y294 (TM2/TM7), and L110.Y294 (TM3/TM7) for the 𝒟_RE_ dataset, and G137.G192 (TM4/TM5), I115.A293 (TM3/TM7), P112.S429 (TM3/TM11) and S234.K471 (TM6/TM12) for the 𝒟_CA_ dataset. Locations of these residue pairs and the corresponding distance histograms are shown in **Figure 9** (for 𝒟_CA_) and **Figure S7** (for 𝒟_RE_).

**Figure 9:**
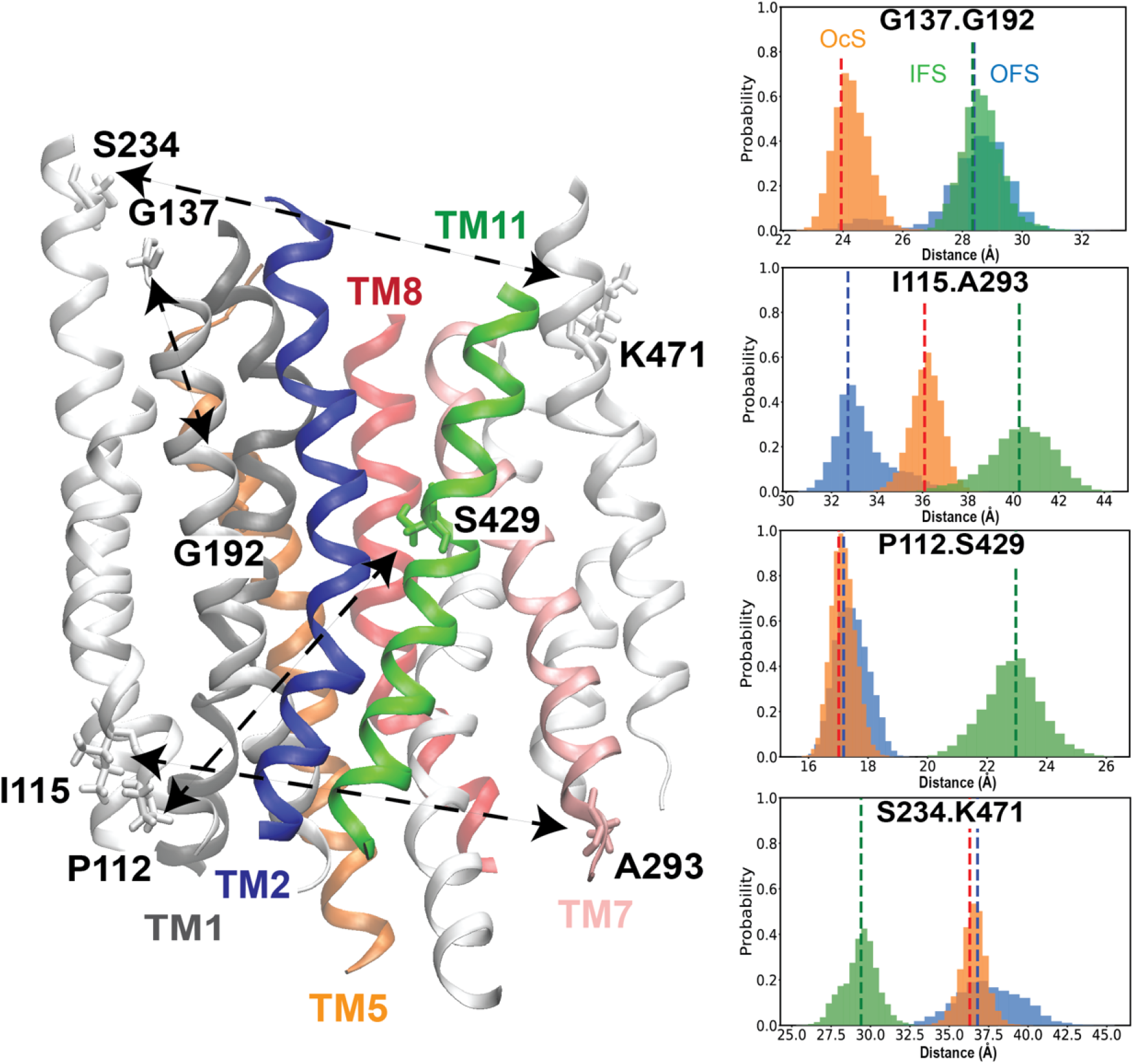
Structural Context for the Key Features derived from the DR analysis of the *𝒟*_CA_ Dataset. The locations of the key features identified by the MPSO-S method are shown on the OcS model of ggMFSD2A. TMs 1, 2, 5, 7, 8, and 11 are color-coded in grey, blue, orange, pink, red, and green, respectively. The residue pairs comprising the key features are shown in licorice and are connected by arrows. Accompanying the structural representation are histograms that show distributions of the distances corresponding to the key features in OFS (in blue), in OcS (in orange), and in IFS (in green).

The histograms illustrate that each feature is effective at distinguishing at least one of the three conformational states of the MFSD2A protein from the others. Specifically, the residue pairs G137.G192 and G137.G250 are key in differentiating the OcS. Conversely, the pairs P112.S429 and S234.K471 were effective in separating the IFS. The G47.I295 distance separates the OFS, and the pairs I115.A293, L81.F308, W87.Y294 and L110.Y294 enable distinguishing all the three states. Thus, positioning of TM7 helix appears to be the most critical structural feature for discriminating OFS, OcS, and IFS, the result that is consistent with our earlier structural and computational studies implicating conformational changes in TM7 during the state-to-state transitions in MFSD2A^30^. These findings underscore the reliability of the identified CVs for separating the functional states in this transporter system.

## DISCUSSION

A growing number of studies address efficient ways of analyzing MD data to extract meaningful structural descriptors, or CVs, that can reliably distinguish different metastable states of a protein system^15,64,80-82^. Knowledge of such descriptors is particularly useful when sampling conformational transitions between these states presents a challenge in conventional MD simulations. Indeed, in such cases, the CVs can guide the design of enhanced sampling simulations in which the state-to-state transitions can be more efficiently sampled.

Our work contributes to the field by proposing a general automated machine learning-based protocol for extracting from MD simulations a minimal yet effective set of CVs that can accurately and efficiently distinguish structurally different states of a molecular system. The automated nature of the protocol reduces the reliance of the CV design process on manual user inspection of MD trajectories and minimizes the need for physical intuition. Here, we tested our protocol on a mechanistically well-studied membrane protein system: the MFSD2A lysolipid transporter. By applying the method to the MD trajectories of MFSD2A in three functionally relevant metastable states – OFS, OcS, and IFS – we demonstrated that the approach enables the automatic detection of CVs that classify the three states with high accuracy and are clearly interpretable in a detailed structural context.

The CVs in our protocol are obtained through a multi-step process which includes MD data filtering, feature selection, and DR. The filtering and feature selection steps eliminate low variance structural features as well as the features that share significant mutual information. The final set of filtered features is combined into CVs via different automated and non-trained linear DR methods. While for the MFSD2A system we utilized pairwise residue distances as features, the protocol is flexible to allow various types of structural inputs (e.g., angular variables, volumetric measures) directly obtained from MD simulations. The choice of input data allows us to generate system-specific CVs, rendering this method versatile for a wide range of systems, provided that the fluctuations in the studied metastable states are sufficiently sampled in MD simulations.

Our analysis systematically examined the impact of various feature selection and DR techniques not only on class separability but also on the number of identified features. Indeed, the increased number of features, while expected to improve class separability, can potentially complicate interpretability of the resulting CVs. We found that, generally, wrapper feature selection methods, such as BPSO and MPSO, outperformed filter selection methods, the χ^2^ test and Fisher scores, in the classification task. Specifically, the χ^2^ test was the least effective, consistently resulting in the poorest class separation, while MPSO showed the highest degree of class separation particularly when used with loose regularization (MPSO-L). However, MPSO-L also yielded the highest number of selected features. At the same time, the performance of MPSO with strong regularization (MPSO-S) was only marginally diminished compared to MPSO-L while the method selected fewer features. Given the comparable performance of MPSO-S and MPSO-L, the identification of fewer features renders MPSO-S the preferred feature selection method for our system. While the wrapper feature selection methods outperformed the filter methods, the latter methods still showed high accuracy. To this end, it is important to note that the χ^2^ test and Fisher score methods are computationally more efficient compared to the PSO methods. Indeed, for our datasets, the filter methods required only seconds to run on 16 CPUs, whereas MPSO, due to its need for hyperparameter optimization and the iterative construction and evaluation of a ML model for each particle in the swarm with each iteration took several minutes per run, resulting in hours of calculations on 48 CPUs. Given these considerations and depending on the available computational resources, the filter methods may be a preferable choice.

For the DR step in our protocol, we compared performance of PCA with various LDA methods. The latter included well-established FLDA, MHLDA, and ZHLDA approaches as well as a new gradient-based LDA variant we proposed, GDHLDA, which integrates the harmonic mean formulations for within-class and between-class variances from ZHLDA and MHLDA. Our results demonstrate that LDA-based methods outperform PCA, showing a higher degree of class separation across all feature selection techniques. Furthermore, we found that all the LDA-based methods follow a highly consistent pattern of feature extraction (i.e., the different LDA methods identify similar structural regions as significant for distinguishing the three metastable states). Interestingly, for the MFSD2A system, we found that the added complexity to the GDHLDA algorithm (i.e., consideration of the harmonic means as well as implementation of the gradient descent optimization) did not yield enhanced performance as the results for GDHLDA approach were similar to those of the other LDA implementations. This can be explained by our finding that the latter LDA approaches already performed class separation task for the MFSD2A system with high accuracy. To better assess potential benefits of the GDHLDA approach, it will be important to compare its performance to all the other LDA methods used here for the molecular systems where structural differences between classes are less evident (in MFSD2A, the IFS is markedly different from both the OcS and OFS, as evidenced by a RMSD of 3.34Å and 3.74Å respectively^29-30^).

To compare how collective movements of various structural segments in MFSD2A and more localized side-chain level structural rearrangements contribute to the classification of the functional states we considered as input features two distinct sets of pairwise distances: 𝒟_RE_, involving distances between the geometric centers of the residues, and 𝒟_CA_, involving distances between the *C*_*α*_ atoms. We found that the algorithm achieved notably better class separation with 𝒟_RE_ dataset. This is expected, as often subtle backbone-level conformational changes in proteins are accompanied by larger changes in the corresponding side-chain distances. One notable example of such behavior in the MFSD2A system is related to global movements of TM7 helix and concomitant local conformational shifts in Y294 residue residing on this helix during transitions between OFS, OcS, and IFS^30^. Thus, while TM7 features were predicted by the algorithm as important for the classification irrespective of the input dataset (i.e., with either 𝒟_CA_ or 𝒟_RE_), Y294 residue was more prevalent when the algorithm used 𝒟_RE_ dataset as input. This suggests that side-chain level conformational shifts in Y294 is a critical feature for classification of the three states. This prediction is consistent with our previous MD simulation studies^30^ in which rotameric changes in Y294 side-chain were found to be mechanistically linked to the state-to-state transitions in MFSD2A.

Overall, the most significant features in the CVs identified by our protocol described changes in the positioning of primarily TMs 2, 5, 7, and 11 (∼73% of the highest contributing features for class separation were associated with these helices), highlighting their critical roles in state-to-state transitions in MFSD2A. This finding is in line with previous structural studies on MFSD2A showing rearrangement of these TM segments during the transitions. Our algorithm further found residues on the outer helices such as TM4 and TM6 as strongly contributing to the CVs (16.3% and 12.7% of the highest contributing features), indicating that the structural changes in the protein during the functional transitions are widespread and extend across the TM bundle, consistent with the proposed rocker switch mechanism for substrate transport by MFSD2A^29-32,^

Interestingly, some feature selection methods highlight structural changes that relate to local helical distortions in the protein. For example, the Fisher method highlights G137-Q198 and G137-I199 distances as significant features. We found that 137-138 and 198-202 regions, both located on the EC ends of TM helices (TM4 and TM5, respectively) undergo folding-unfolding transitions as MFSD2A switches between the three states. Given their terminal locations, it is not clear whether the observed helical distortions are important part of the state-to-state transition mechanism in MFSD2A. Indeed, the alternative feature selection strategies did not detect these helical distortions as important features. Nevertheless, the ability of our protocol to register these local dynamic modes can be useful for molecular systems in which such helical distortions are essential part of the protein functional mechanism.

The overall performance of our LDA-based classification models was evaluated by comparing their predictions to those obtained from the supervised non-linear DCNN algorithm for pattern recognition analysis of MD trajectories we developed and implemented earlier^36^. The DCNN identified TM6 and TM7 helices as the most important for classifying the metastable states in MFSD2A, the regions also predicted by the LDA methods. This consistency underscores the robustness of the LDA approaches for class separation. Importantly, the automated nature of the LDA algorithm (i.e., no data training is required) and its high interpretability suggest that the LDA methods can be effective when incorporated into a pipeline for analyzing MD simulations of complex molecular systems for CV detection.

While the protocol described here is efficient and automated, it needs to be noted that depending on the considerations discussed above (specific feature selection and/or DR approach used) there could be some variations in the identified CVs. Therefore, the final decision on which set of CVs are the most optimal in separating classes may not be obvious. Several potential solutions to this problem include: 1) selecting the CV set that showed the highest degree of class separation (however, this may result in selecting relatively large number of CVs); 2) selecting the CVs that were the most commonly predicted by the protocol across all the feature selection and DR methods used; 3) combining CVs from both 𝒟_CA_ and 𝒟_RE_ datasets obtained from the same feature selection and DR methods. The ultimate test and validation of the identified CVs should come in future studies where the CVs would be used to drive biased MD simulations. Such simulations will demonstrate the ability of the selected CVs to sample efficiently conformational transitions in the protein.

## AUTHOR CONTRIBUTIONS

MO, MR and GK designed the research; MR carried out MD simulations; MO, MR, and HX carried out the data analyses; all the authors discussed the results and wrote the manuscript.

## ACKNOWLEDGMENTS

GK gratefully acknowledges support from the 1923 Fund. MO is supported by Lili Foo Hing fellowship. MR gratefully acknowledges support from Weill Cornell Graduate School “Markey Graduate School of Medical Science Fellowship”. The authors are grateful for the computational resources under Project BIP109 at the Oak Ridge Leadership Computing Facility, which is a DOE Office of Science User Facility supported under Contract DE-AC05-00OR22725, and for the in-house computational resources of the David A. Cofrin Center for Biomedical Information in the Institute for Computational Biomedicine at Weill Cornell Medical College.

## SUPPLEMENTAL MATERIALS

**Figure S1.**
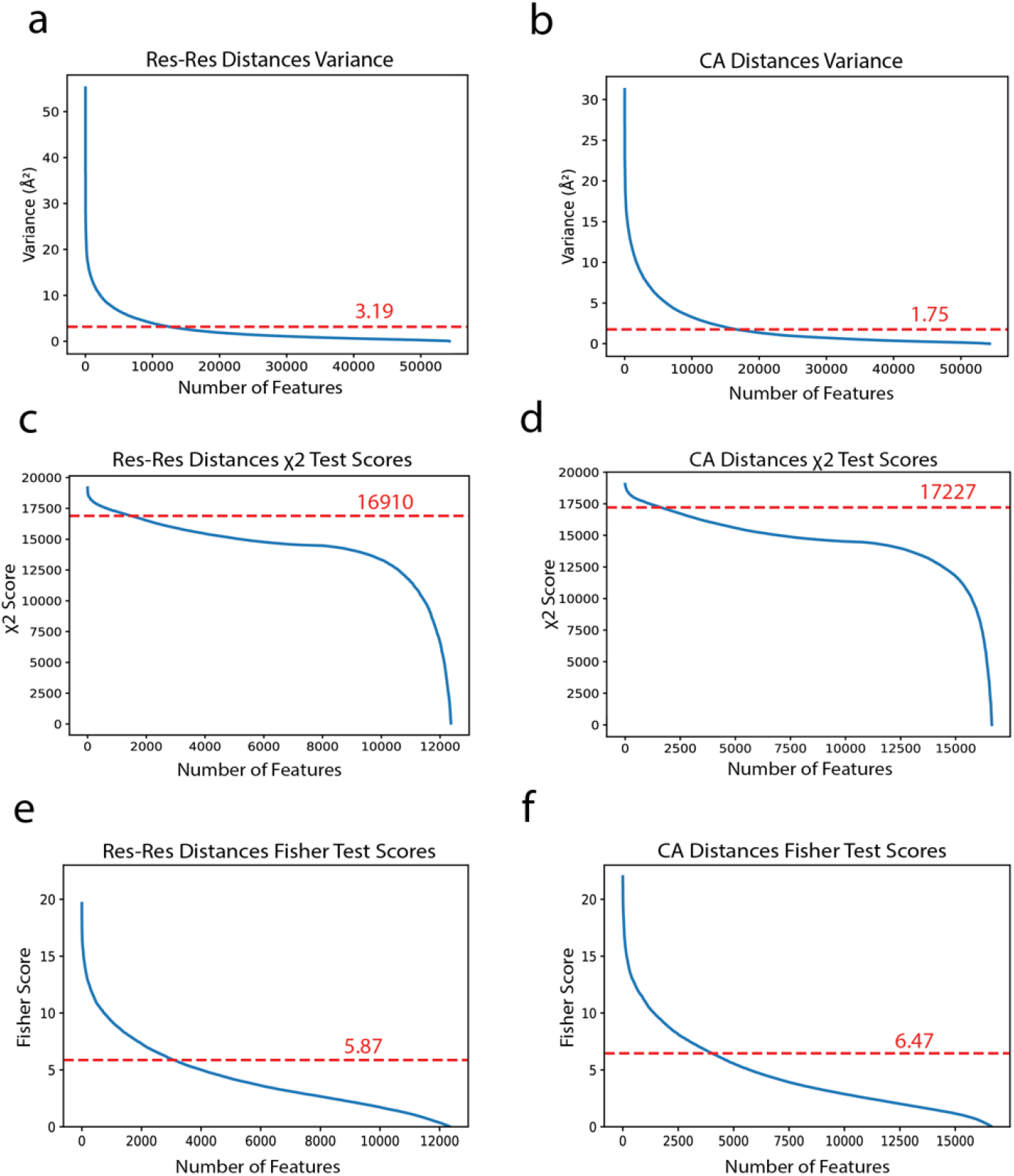
Elbow Point Analysis for Threshold Identification. **(a-b)** Variance plots for **(a)** 𝒟_RE_ and **(b)** 𝒟_CA_ pairwise distances used as features. **(c-d)** χ^2^ test score plots for **(c)** 𝒟_RE_ and **(d)** 𝒟_CA_ pairwise distances. **(e-f)** Fisher score plots for **(e)** 𝒟_RE_ and **(f)** 𝒟_CA_ pairwise distances. All plots illustrate a sharp decline, reaching a plateau which signifies the threshold for feature selection based on the Kneedle algorithm.

**Figure S2.**
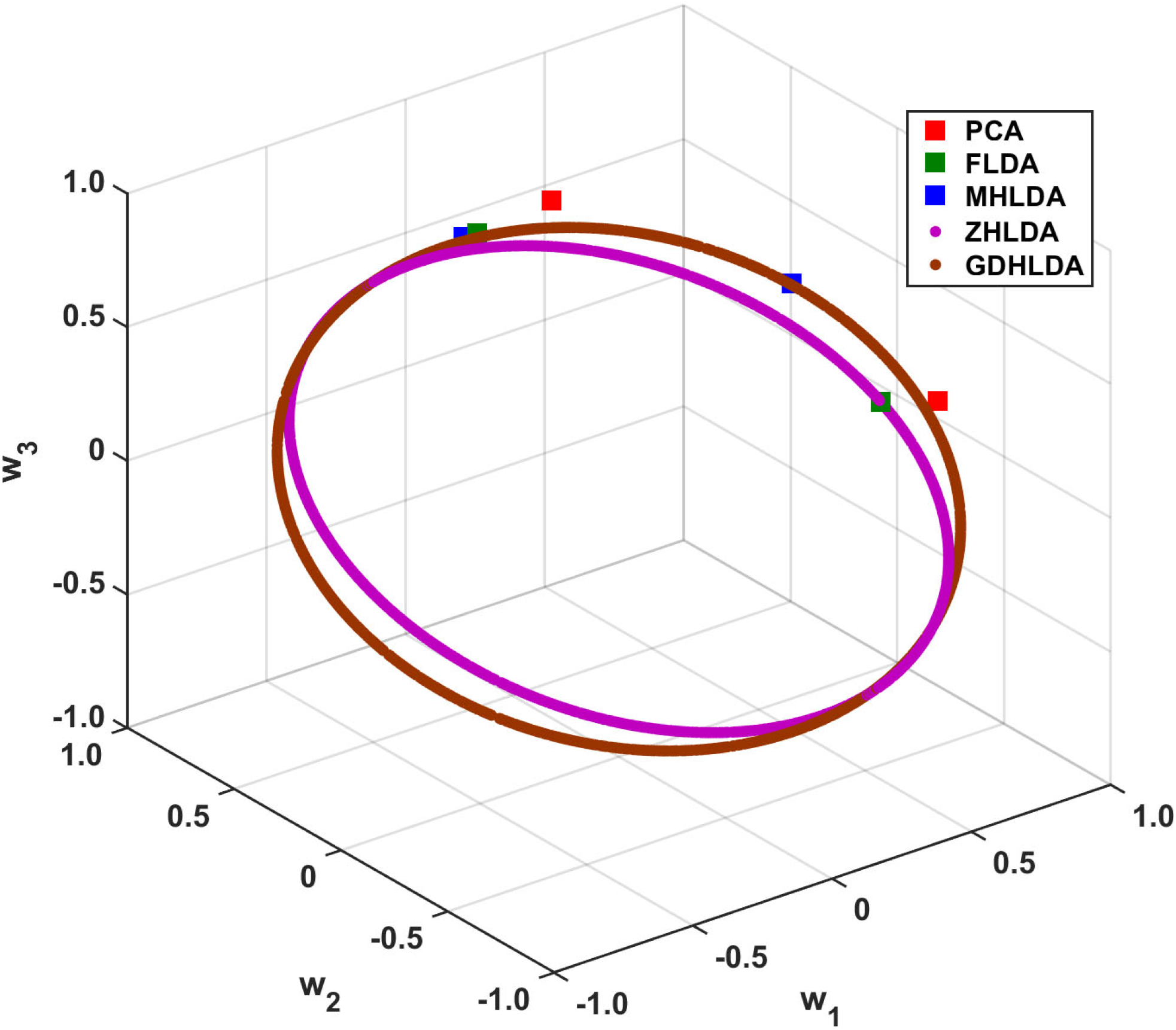
Visualization of solution spaces defined by eigenvectors from eigendecomposition-based methods and gradient descent-based methods. Every eigenvector is represented as a distinct point, with its position determined by the corresponding feature weights *w*_*i*_. The coloring scheme is the same as in Figures S2 and S3.

**Figure S3:**
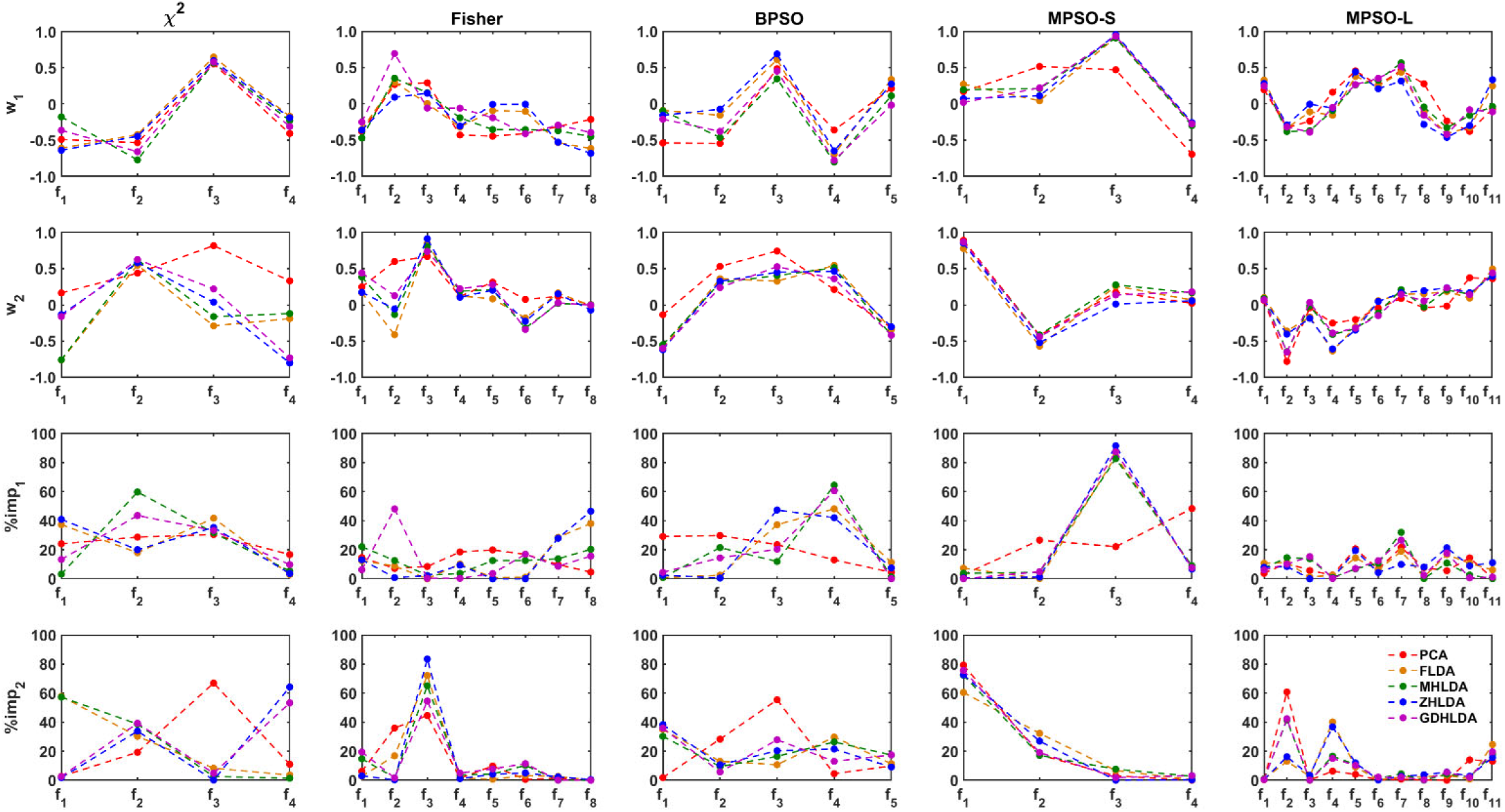
Plots of feature weights and importance scores for the *𝒟*_*CA*_ dataset. The first row displays the weights *w*_1_ of features (selected by χ^2^, Fisher, BPSO, MPSO-S, and MPSO-L) in CV1, as calculated by PCA (red), FLDA (orange), MHLDA (green), ZHLDA (blue), and GDHLDA (purple). The second row presents the weights *w*_2_ for CV2 using the same DR methods and coloring scheme. The third and fourth rows show the importance scores (%imp_1_ and %imp_2_) of features in CV1 and CV2, respectively, using the same DR methods and color coding. *f*_*i*_ represents the *i* ^th^ feature in the cell of Table 2 corresponding to the given combination of the feature selection and DR methods.

**Figure S4:**
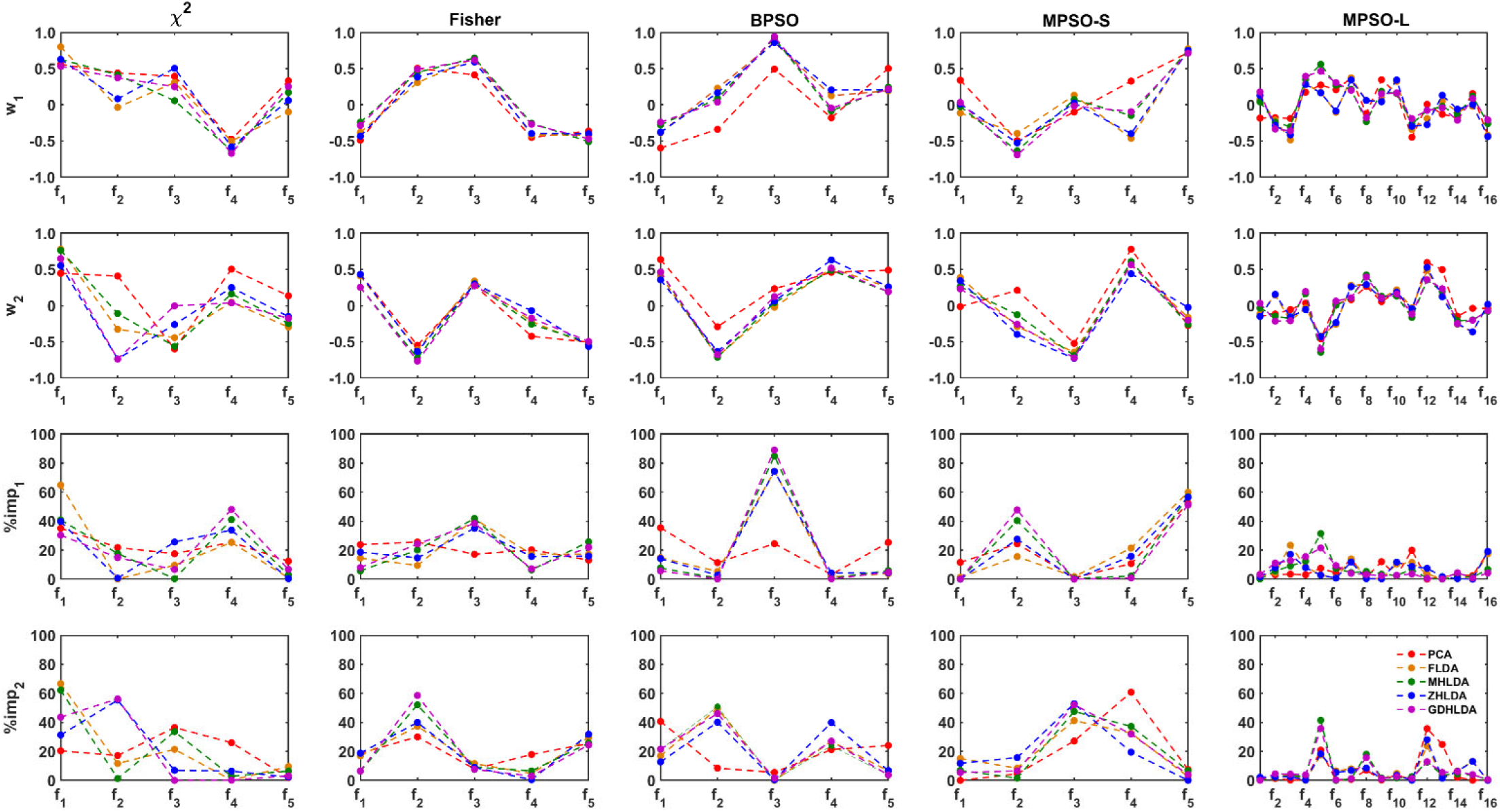
Plots of feature weights and importance scores for the *𝒟*_RE*S*_ dataset. The first row displays the weights *w*_1_ of features (selected by χ^2^, Fisher, BPSO, MPSO-S, and MPSO-L) in CV1, as calculated by PCA (red), FLDA (orange), MHLDA (green), ZHLDA (blue), and GDHLDA (purple). The second row presents the weights *w*_2_ for CV2 using the same DR methods and coloring scheme. The third and fourth rows show the importance scores (%imp_1_ and %imp_2_) of features in CV1 and CV2, respectively, using the same DR methods and color coding. *f*_*i*_ represents the *i* ^th^ feature in the cell of Table 2 corresponding to the given combination of the feature selection and DR methods.

**Figure S5:**
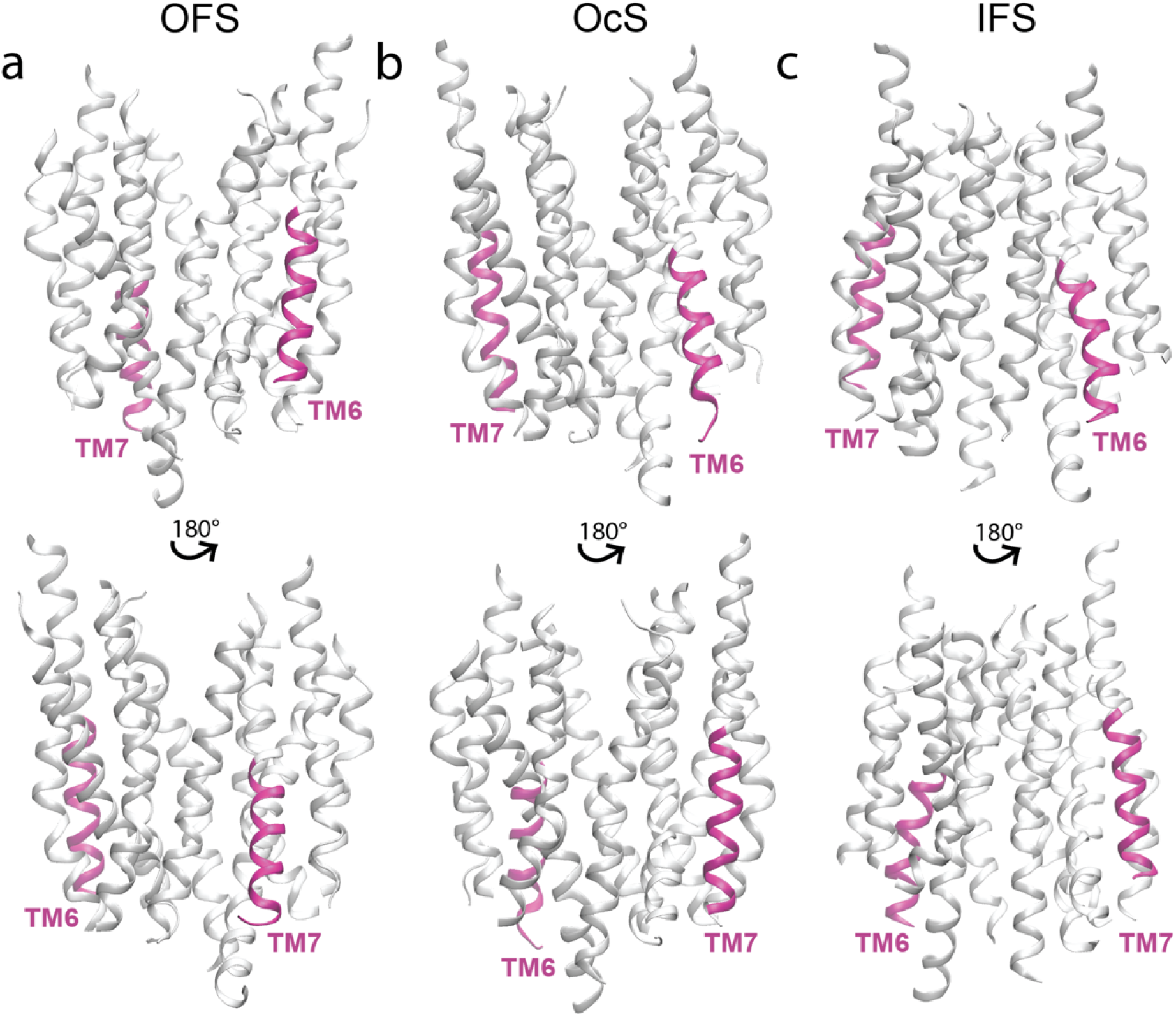
Structural motifs on ggMFSD2A predicted by the DCNN algorithm as the most significant for classification. **(a)** OFS **(b)** OcS and **(c)** IFS structures of ggMFSD2A highlighting the locations of the most significant residues for classification based on the saliency analysis. In each panel, the protein is depicted in two orientations differing by a 180-degree rotation as indicated. The most salient features, the IC end of TM6 (residues 249 to 263), and the IC end of TM7 (residues 292 to 306), are colored in pink, while the rest of the protein is show in grey.

**Figure S6:**
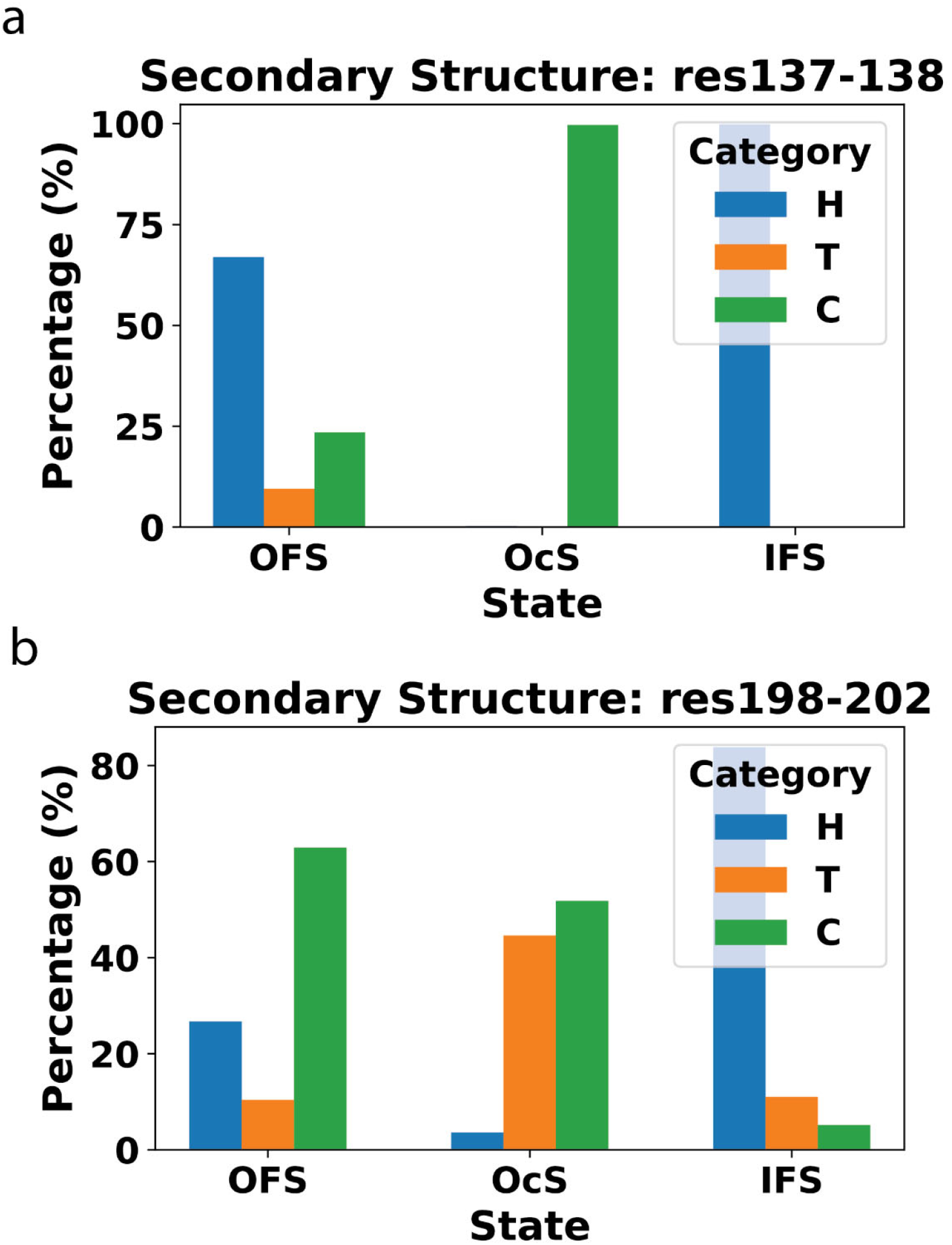
Secondary Structure Analysis of ggMFSD2A: The bar graphs display the percentage of three secondary structure categories (H: Helix, T: Turn, C: Coil) at two different regions, residues 137-138 (a) and 198-202 (b) of ggMFSD2A in OFS, OcS and IFS. The results show: 1) a predominant coil structure for the 137-138 region in the OcS, suggesting unwinding from the OFS and IFS, where the helix structure is more prevalent. 2) a decrease in helical structure in the 198-202 region when transitioning from IFS to OFS and OcS.

**Figure S7:**
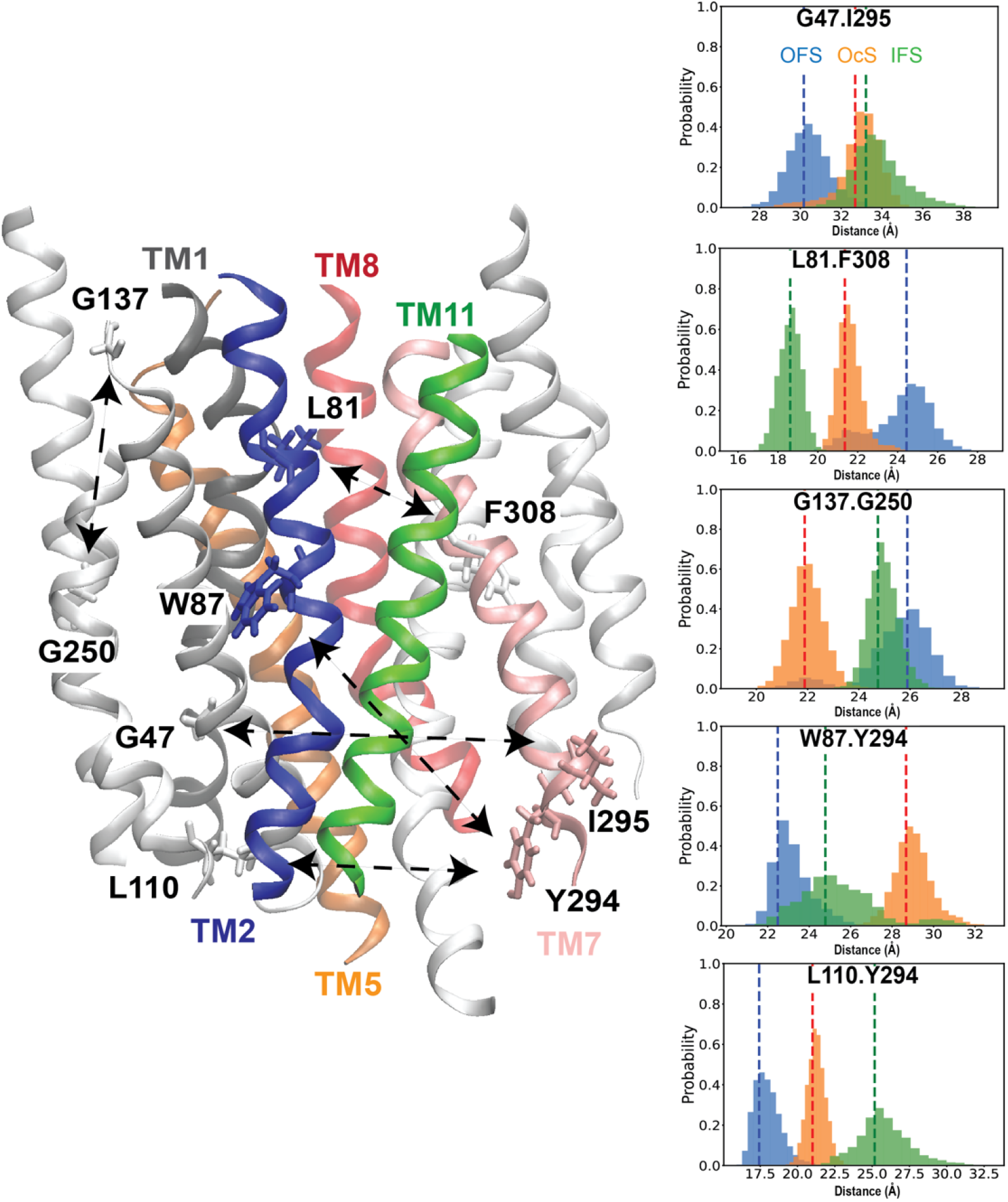
Structural Context for the Key Features derived from the DR analysis of *𝒟*_RES_ Dataset. The locations of the key features identified by the MPSO-S method are shown on the OcS model of ggMFSD2A. TMs 1, 2, 5, 7, 8, and 11 are color-coded in grey, blue, orange, pink, red, and green, respectively. The residue pairs comprising the key features are shown in licorice and are connected by arrows. Accompanying the structural representation are histograms that show distributions of the distances corresponding to the key features in OFS (in blue), in OcS (in orange), and in IFS (in green).

**Table S1:**
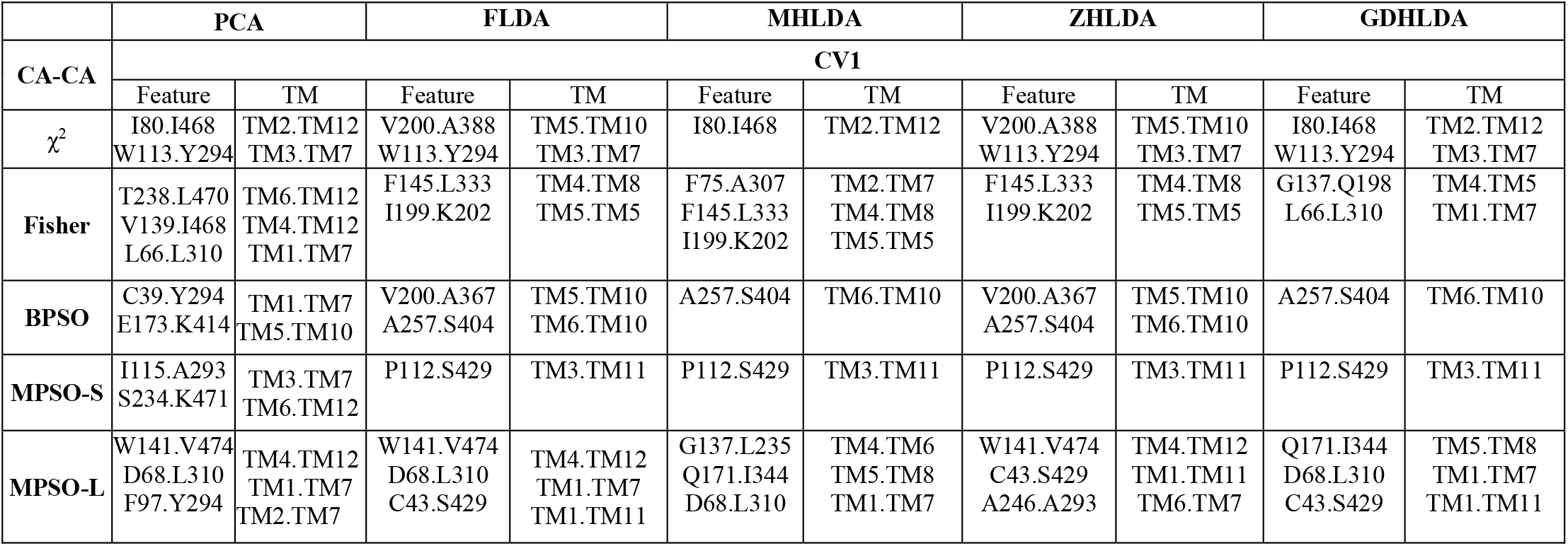
Summary of the most significant features for CV1 in the *𝒟*_CA_ dataset. Each row displays the features with importance scores (from Fig. S3) totaling more than 50% selected by different feature selection methods (χ^2^, Fisher, BPSO, MPSO-S, MPSO-L) for CV 1 calculated using different DR methods (PCA, FLDA, ZHLDA, MHLDA, GDHLDA).

|  | PCA |  | FLDA |  | MHLDA |  | ZHLDA |  | GDHLDA |  |
| --- | --- | --- | --- | --- | --- | --- | --- | --- | --- | --- |
| CA-CA | CV1 |  |  |  |  |  |  |  |  |  |
|  | Feature | TM | Feature | TM | Feature | TM | Feature | TM | Feature | TM |
| $\chi^2$ | I80.I468<br>W113.Y294 | TM2.TM12<br>TM3.TM7 | V200.A388<br>W113.Y294 | TM5.TM10<br>TM3.TM7 | I80.I468 | TM2.TM12 | V200.A388<br>W113.Y294 | TM5.TM10<br>TM3.TM7 | I80.I468<br>W113.Y294 | TM2.TM12<br>TM3.TM7 |
| Fisher | T238.L470<br>V139.I468<br>L66.L310 | TM6.TM12<br>TM4.TM12<br>TM1.TM7 | F145.L333<br>I199.K202 | TM4.TM8<br>TM5.TM5 | F75.A307<br>F145.L333<br>I199.K202 | TM2.TM7<br>TM4.TM8<br>TM5.TM5 | F145.L333<br>I199.K202 | TM4.TM8<br>TM5.TM5 | G137.Q198<br>L66.L310 | TM4.TM5<br>TM1.TM7 |
| BPSO | C39.Y294<br>E173.K414 | TM1.TM7<br>TM5.TM10 | V200.A367<br>A257.S404 | TM5.TM10<br>TM6.TM10 | A257.S404 | TM6.TM10 | V200.A367<br>A257.S404 | TM5.TM10<br>TM6.TM10 | A257.S404 | TM6.TM10 |
| MPSO-S | I115.A293<br>S234.K471 | TM3.TM7<br>TM6.TM12 | P112.S429 | TM3.TM11 | P112.S429 | TM3.TM11 | P112.S429 | TM3.TM11 | P112.S429 | TM3.TM11 |
| MPSO-L | W141.V474<br>D68.L310<br>F97.Y294 | TM4.TM12<br>TM1.TM7<br>TM2.TM7 | W141.V474<br>D68.L310<br>C43.S429 | TM4.TM12<br>TM1.TM7<br>TM1.TM11 | G137.L235<br>Q171.I344<br>D68.L310 | TM4.TM6<br>TM5.TM8<br>TM1.TM7 | W141.V474<br>C43.S429<br>A246.A293 | TM4.TM12<br>TM1.TM11<br>TM6.TM7 | Q171.I344<br>D68.L310<br>C43.S429 | TM5.TM8<br>TM1.TM7<br>TM1.TM11 |

**Table S2:** Summary of the most significant features for CV2 in the *𝒟*_CA_ dataset. Each row displays the features with importance scores (from Fig. S3) totaling more than 50% selected by different feature selection methods (χ^2^, Fisher, BPSO, MPSO-S, MPSO-L) for CV 2 calculated using different DR methods (PCA, FLDA, ZHLDA, MHLDA, GDHLDA).

|  | PCA |  | FLDA |  | MHLDA |  | ZHLDA |  | GDHLDA |  |
| --- | --- | --- | --- | --- | --- | --- | --- | --- | --- | --- |
| CA-CA | CV2 |  |  |  |  |  |  |  |  |  |
|  | Feature | TM | Feature | TM | Feature | TM | Feature | TM | Feature | TM |
| $\chi^2$ | W113.Y294 | TM3.TM7 | V200.A388 | TM5.TM10 | V200.A388 | TM5.TM10 | L81.G392 | TM2.TM10 | L81.G392 | TM2.TM10 |
| Fisher | G137.Q198<br>G137.I199 | TM4.TM5<br>TM4.TM5 | G137.I199 | TM4.TM5 | G137.I199 | TM4.TM5 | G137.I199 | TM4.TM5 | G137.I199 | TM4.TM5 |
| BPSO | V200.A367 | TM5.TM10 | C39.Y294<br>A257.S404 | TM1.TM7<br>TM6.TM10 | C39.Y294<br>A257.S404 | TM1.TM7<br>TM6.TM10 | C39.Y294<br>A257.S404 | TM1.TM7<br>TM6.TM10 | C39.Y294<br>V200.A367 | TM1.TM7<br>TM5.TM10 |
| MPSO-S | G137.G192 | TM4.TM5 | G137.G192 | TM4.TM5 | G137.G192 | TM4.TM5 | G137.G192 | TM4.TM5 | G137.G192 | TM4.TM5 |
| MPSO-L | G137.L235 | TM4.TM6 | V200.T304<br>A246.A293 | TM5.TM7<br>TM6.TM7 | G137.L235<br>A246.A293 | TM4.TM6<br>TM6.TM7 | G137.L235<br>V200.T304 | TM4.TM6<br>TM5.TM7 | G137.L235<br>A246.A293 | TM4.TM6<br>TM6.TM7 |

**Table S3:**
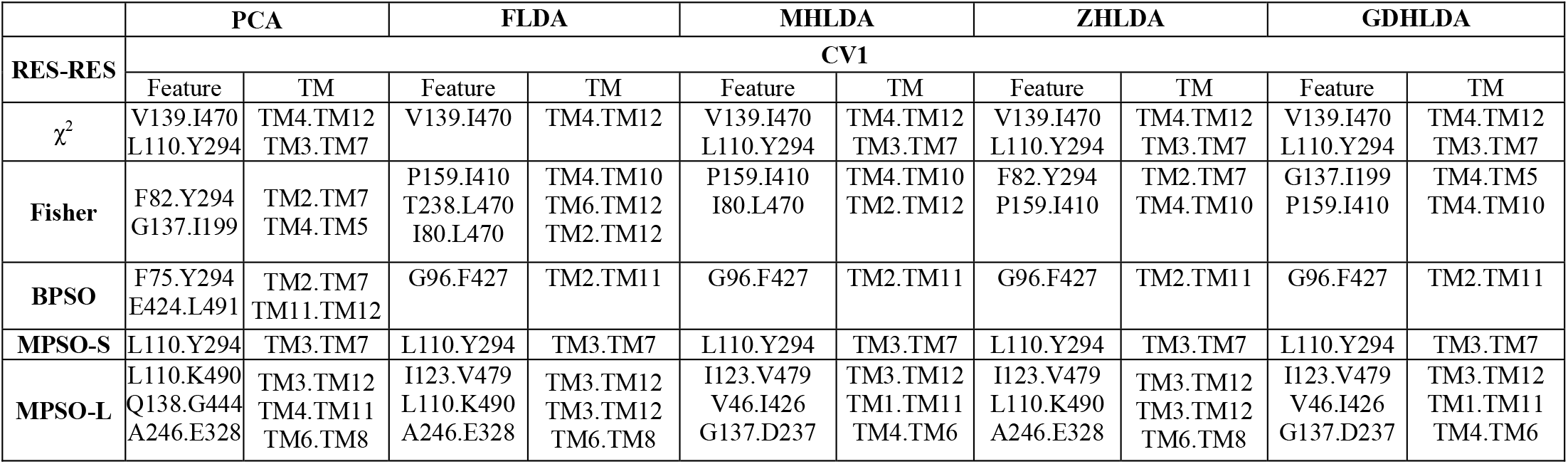
Summary of the most significant features for CV1 in the *𝒟*_RES_ dataset. Each row displays the features with importance scores (from Fig. S4) totaling more than 50% selected by different feature selection methods (χ^2^, Fisher, BPSO, MPSO-S, MPSO-L) for CV 1 calculated using different DR methods (PCA, FLDA, ZHLDA, MHLDA, GDHLDA).

**Table S4:** Summary of the most significant features for CV2 in the *𝒟*_**RES**_ dataset. Each row displays the features with importance scores (from Fig. S4) totaling more than 50% selected by different feature selection methods (χ^2^, Fisher, BPSO, MPSO-S, MPSO-L) for CV 2 calculated using different DR methods (PCA, FLDA, ZHLDA, MHLDA, GDHLDA).

|  | PCA |  | FLDA |  | MHLDA |  | ZHLDA |  | GDHLDA |  |
| --- | --- | --- | --- | --- | --- | --- | --- | --- | --- | --- |
| RES-RES | CV2 |  |  |  |  |  |  |  |  |  |
|  | Feature | TM | Feature | TM | Feature | TM | Feature | TM | Feature | TM |
| $\chi^2$ | V200.A388<br>L110.Y294 | TM5.TM10<br>TM3.TM7 | V139.I470 | TM4.TM12 | V139.I470 | TM4.TM12 | I80.I468 | TM2.TM12 | I80.I468 | TM2.TM12 |
| Fisher | G137.I199<br>I80.L470 | TM4.TM5<br>TM2.TM12 | G137.I199<br>I80.L470 | TM4.TM5<br>TM2.TM12 | G137.I199 | TM4.TM5 | F82.Y294<br>G137.I199 | TM2.TM7<br>TM4.TM5 | G137.I199 | TM4.TM5 |
| BPSO | F75.Y294<br>E424.L491 | TM2.TM7<br>TM11.TM12 | F75.S305 | TM2.TM7 | F75.S305 | TM2.TM7 | F75.S305<br>Y294.A391 | TM2.TM7<br>TM7.TM10 | F75.S305<br>Y294.A391 | TM2.TM7<br>TM7.TM10 |
| MPSO-S | W87.Y294 | TM2.TM7 | G137.G250<br>W87.Y294 | TM4.TM6<br>TM2.TM7 | G137.G250<br>W87.Y294 | TM4.TM6<br>TM2.TM7 | G137.G250 | TM4.TM6 | G137.G250 | TM4.TM6 |
| MPSO-L | F149.Y294<br>Y294.L333 | TM5.TM7<br>TM7.TM8 | G137.D237<br>F149.Y294<br>R181.L415 | TM4.TM6<br>TM5.TM7<br>TM5.TM10 | G137.D237<br>Q138.A241 | TM4.TM6<br>TM4.TM6 | G137.D237<br>F149.Y294<br>R181.L415 | TM4.TM6<br>TM5.TM7<br>TM5.TM10 | G137.D237<br>Q138.A241 | TM4.TM6<br>TM4.TM6 |

### Geometrical Analysis on the Solution Spaces Given by the Linear DR Algorithms

Unlike PCA, LDA methods such as FLDA and MHLDA do not generally guarantee orthogonality of eigenvectors (or CVs) since the product 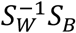 is not necessarily symmetric. Additionally, solving the generalized eigenvalue problem, which involves the eigendecomposition of 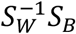, yields a single pair of eigenvectors. This scenario, however, does not capture the non-uniqueness of the basis set of the subspace, as theoretically, an infinite number of basis vectors can span the same subspace. The gradient descent-based methods, ZHLDA and GDHLDA, address this limitation by capturing all possible orthogonal basis vectors that define the subspace, provided the gradient descent algorithm initiates from varied starting points. Consequently, we hypothesize that in a solution space where the eigenvector is plotted against the weights of the features, a unit circle centered at the origin should emerge (owing to the normalization of eigenvectors), with its orientation in the solution space being determined by different averaging schemes (arithmetic or harmonic averaging for within-class and between-class scatter matrices). To enhance both visualization and understanding, we opted for three specific features (*f*_1_: C39:Y294, *f*_3_: V200:A367, *f*_4_: A257:S404) chosen by BPSO-SVM from the dataset 𝒟_RE_ for generating CVs as linear combinations of these features using both ZHLDA and GDHLDA approaches. Subsequently, we initiated the gradient descent algorithms from various starting points. As illustrated in **Figure S2**, the solution space was found to be defined by a unit circle (colored in red for ZHLDA and green for GDHLDA) centered at the origin, with its orientation contingent upon the applied averaging scheme.

